# The psychosis risk factor *RBM12* encodes a novel repressor of GPCR/cAMP signal transduction

**DOI:** 10.1101/2023.01.12.523776

**Authors:** Khairunnisa M. Semesta, Angelica Garces, Nikoleta G. Tsvetanova

## Abstract

*RBM12* is a high-penetrance risk factor for familial schizophrenia and psychosis, yet its precise cellular functions and the pathways to which it belongs are not known. We utilize two complementary models, HEK293 cells and human iPSC-derived neurons, and delineate RBM12 as a novel repressor of the G protein-coupled receptor/cyclic AMP/protein kinase A (GPCR/cAMP/PKA) signaling axis. We establish that loss of RBM12 leads to hyperactive cAMP production and increased PKA activity as well as altered neuronal transcriptional responses to GPCR stimulation. Notably, the cAMP and transcriptional signaling steps are subject to discrete RBM12-dependent regulation. We further demonstrate that the two *RBM12* truncating variants linked to familial psychosis impact this interplay, as the mutants fail to rescue GPCR/cAMP signaling hyperactivity in cells depleted of RBM12. Lastly, we present a mechanism underlying the impaired signaling phenotypes. In agreement with its activity as an RNA-binding protein, loss of RBM12 leads to altered gene expression, including that of multiple effectors of established significance within the receptor pathway. Specifically, the abundance of adenylyl cyclases, phosphodiesterase isoforms, and PKA regulatory and catalytic subunits is impacted by RBM12 depletion. We note that these expression changes are fully consistent with the entire gamut of hyperactive signaling outputs. In summary, the current study identifies a previously unappreciated role for RBM12 in the context of the GPCR/cAMP pathway that could be explored further as a tentative molecular mechanism underlying the functions of this factor in neuronal physiology and pathophysiology.

## Introduction

G protein-coupled receptors (GPCRs) mediate essential aspects of human physiology and make up the targets of more than a third of all clinically prescribed drugs.^1^ Because of their vast physiological roles and pharmacological significance, it has been a long-standing goal to identify the factors that regulate GPCR function as these may represent improved targets for therapeutic intervention. Indeed, the main regulatory steps of the cascade are well-characterized. Upon binding to agonist, the receptor stimulates its associated heterotrimeric G protein complex to initiate cell signaling, often via generation of second messengers. Stimulatory receptors, which comprise a large fraction of the GPCR family, couple to Gαs to activate adenylyl cyclases and lead to production of cyclic AMP (cAMP). The GPCR is then ‘shut off’ following phosphorylation by G protein-coupled receptor kinases (GRKs) and engagement of arrestins.^2^ The propagation of the cAMP signal, in turn, is controlled by the interplay between phosphodiesterase (PDE) and effector binding. Protein kinase A (PKA) is one of the main effectors activated by cAMP, and it phosphorylates a myriad of cellular substrates, including the nuclear transcription factor CREB, which stimulates gene transcription.^3^ Yet, despite these significant advances, much remains to be learned about the mechanisms mediating GPCR function. This has been underscored by the recent discovery that GPCR signaling is compartmentalized through unknown mechanisms,^4–7^ which has opened new avenues for exploration.

Through a genome-wide CRISPR screen, we identified close to 100 novel regulators of transcriptional signaling downstream of the prototypical Gαs-coupled beta-2-adrenergic receptor (β2-AR).^8^ From this screen, RNA-binding motif 12 (RBM12) emerged as one of the most prominent candidate-novel repressor of β2-AR/cAMP signaling. Interestingly, RBM12 has no known ties to GPCR biology. It is annotated as an RNA-binding protein (RBP) based on co-immunoprecipitation with RNA^9^ and based on its amino acid sequence, which is predicted to contain three RNA-recognition motifs (RRMs).^10^ In terms of physiology, RBM12 was recently discovered as a high-penetrance risk factor for familial schizophrenia and psychosis in a family-based whole-genome study to identify rare coding-sequence variants associated with disease segregation in a pedigree.^11^ In addition, a different set of mutations in the mouse *RBM12* gene led to neurodevelopmental defects characterized by open mid and forebrain.^12^ Outside of its functions in the brain, RBM12 was found to be a repressor of fetal hemoglobin expression.^9^ Therefore, RBM12 is of clear significance for a range of physiologies. Yet, the cellular functions and pathways to which RBM12 belongs are unknown, and therefore it remains unclear how mutations in this gene contribute to disease pathobiology.

In this study, we report a novel mechanism that ties RBM12 to the GPCR/cAMP signaling cascade. In two complementary models, HEK293 cells and human neurons derived from induced pluripotent stem cells (hiPSCs), we demonstrate that loss of RBM12 leads to hyperactive cAMP production, increased PKA activity, and supraphysiological transcriptional signaling in response to β2-AR stimulation. As a result, the compartmentalization of the β2 - AR/cAMP signal is compromised. Further, the full repertoire of β2-AR-dependent neuronal transcriptional responses is disrupted by loss of RBM12, leading to overall increase in the extent of gene induction for shared target mRNAs as well as to transcription of distinct targets. We also find that the two *RBM12* truncating mutations linked to familial psychosis are likely loss-of-function, as the mutant proteins fail to rescue GPCR/cAMP signaling hyperactivity in cells depleted of RBM12. Lastly, we present an underlying mechanism for these phenotypes by showing that loss of RBM12 leads to increased expression of multiple adenylyl cyclases and the PKA catalytic subunits, and decreased expression of PDE isoforms and a PKA regulatory subunit. In summary, we identify a previously unappreciated role for RBM12 as a repressor of the GPCR/PKA/cAMP signaling axis that is conserved across cell model systems.

## Results

### RBM12 is a novel repressor of GPCR/cAMP signaling

In our published genome-wide CRISPR screen, we employed sorting of gRNA-transduced cells expressing a fluorescent CREB transcriptional reporter following cAMP stimulation through the β2-AR.^8^ We reanalyzed these data ranking the genes based on “hit strength”, defined as the product of the gRNA read count ratio in high versus low fluorescent reporter-expressing sorted fractions and the respective significance *p*-value. With this ranking, RBM12 stood out among the hits, since its depletion gave rise to one of the strongest phenotypes amongst the candidate novel repressors of the pathway (Fig. 1a).

**Figure 1.**
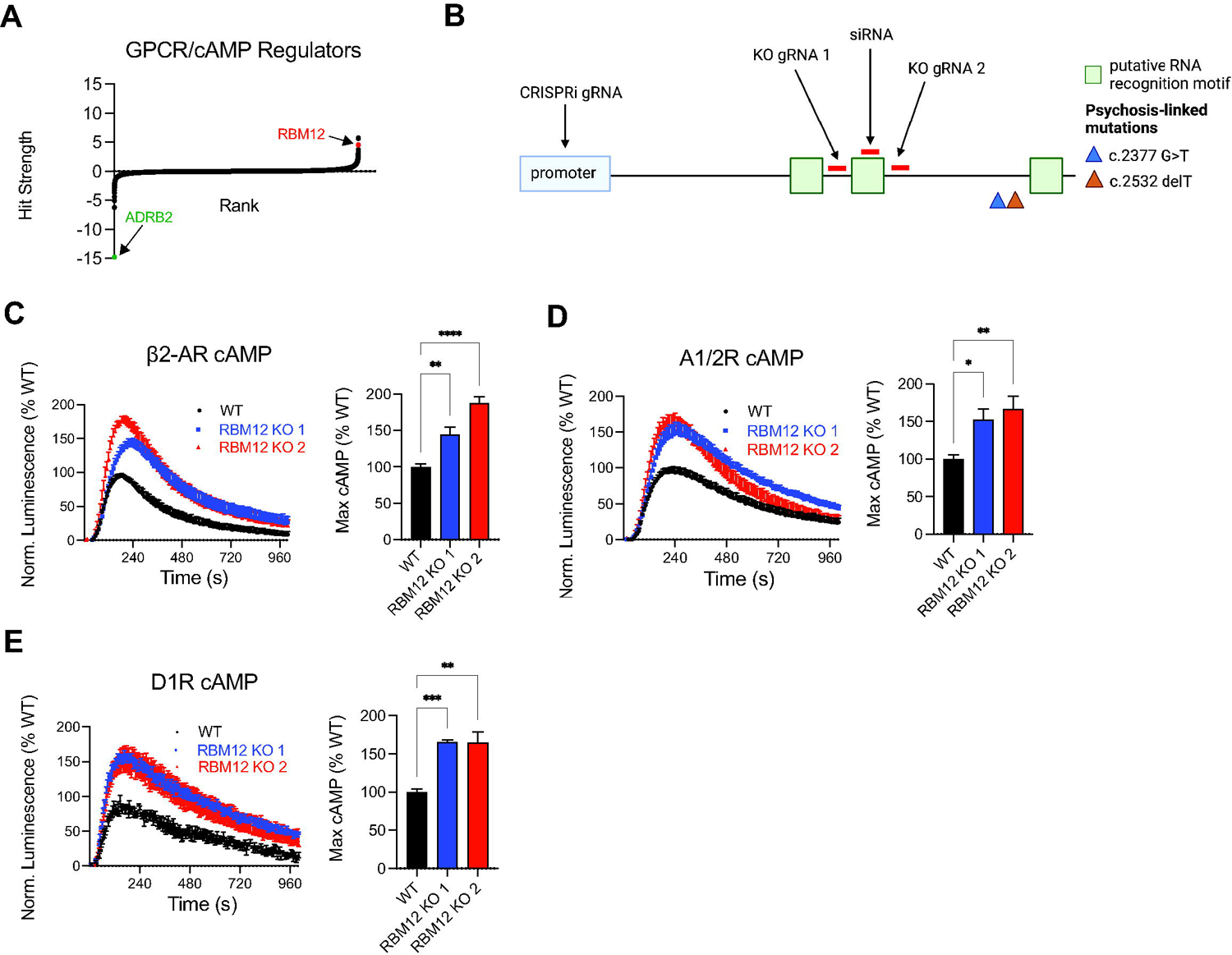
RBM12 loss leads to hyperactive GPCR/cAMP signaling. (A) Genes (20,528) ordered by hit strength (product of phenotype score and -log10(*p*-value)). Candidate repressors exhibit hit strength > 0, and candidate activators exhibit hit strength < 0 in a genome-wide CRISPR screen for GPCR/cAMP regulators.^8^ (B) Schematic of multiple strategies to deplete RBM12 in HEK293 using CRISPRi, RNAi, and CRISPR knockout, depicting positions of guide RNAs, siRNA, and *RBM12* SNPs (c.2377G>T and c.2532delT) implicated in psychosis. (C) Luminescent GloSensor measurement of cAMP accumulation in response to β2-AR agonist isoproterenol (Iso), 100 nM (n = 8). (D) Luminescent GloSensor measurement of cAMP accumulation in response to activation of A1/2R with 10 μM NECA (n = 8). (E) Luminescent GloSensor measurement of cAMP accumulation in response to D1R-selective agonist SKF-81297, 10 nM (n = 4). All data are mean ± SEM. Statistical significance was determined using one-way ANOVA with Dunnett’s correction. See also Figure S1.

To dissect the functions of RBM12 within the GPCR pathway, we generated clonal knockout HEK293 cell lines with CRISPR/Cas9 and two independent gRNAs (Fig. 1b). Characterization of the clonal lines showed that they harbored different frameshift mutations resulting in truncated RBM12 protein missing either one (KO 1) or two (KO 2) of its putative RRM domains (Fig. S1a-b). β2-AR couples to Gαs to stimulate cAMP production, and therefore we first focused on the effect of RBM12 depletion on cAMP generation. Using a cytosolic luciferase-based biosensor to detect cAMP accumulation in real time in intact cells,^13^ we found that β2-AR activation led to hyperactive cytosolic cAMP signaling in the two knockout lines compared to wild-type parental cells (Fig. 1c). We then asked whether the regulatory role of RBM12 is selective for the β2-AR pathway by examining cAMP accumulation downstream of other Gαs-coupled receptors. We observed that stimulation of endogenously expressed adenosine receptors and transfected dopamine 1 receptors led to similarly increased cAMP production in RBM12 knockout cells (Fig. 1d-e and Fig. S1c). In contrast, loss of RBM12 did not have a reproducible impact on cAMP inhibition by Gαi-coupled receptors. Specifically, we surveyed the dopamine 2, delta- and mu-opioid receptors, three prototypical Gαi-coupled GPCRs with important neurobiological functions.^14,15^ We found that the inhibitory activity of these receptors did not differ between wild-type and knockout cells (Fig. S1d-f). Together, these results support RBM12 as a bona fide novel repressor of the GPCR/cAMP pathway and pinpoint a broader regulatory role that extends to multiple GPCRs coupled to the stimulatory G protein.

### Loss of RBM12 leads to increased PKA activity, supraphysiological transcriptional responses, and impaired compartmentalization of β2-AR signaling

PKA activation takes place downstream of cAMP production and mediates CREB phosphorylation and CREB-dependent transcriptional signaling. Therefore, to gain a more complete understanding of the functional effects of loss of RBM12 on GPCR/cAMP signaling, we next examined PKA activity and gene expression changes downstream of β2-AR stimulation.

We began by transfecting wild-type and knockout cells with the single-fluorophore excitation PKA biosensor, ExRai-AKAR2. This biosensor undergoes a conformational change upon PKA-dependent phosphorylation leading to increased fluorescence intensity.^16^ Using microscopy, we detected robust kinase activity in response to isoproterenol across all cell lines. However, RBM12 knockouts displayed higher ExRai-AKAR2 activity in comparison to wild-type (ΔF/F_max_= 1.09 ± 0.12 and 0.82 ± 0.06 in the depleted lines versus 0.60 ± 0.07 in the parental line) (Fig. 2a). To evaluate the impact on transcriptional signaling, we utilized two complementary assays. First, we quantified accumulation of a fluorescent CREB transcriptional reporter^8^ using flow cytometry, and found increased β2-AR-dependent reporter accumulation in the knockouts (Fig. 2b). Next, we used RT-qPCR analysis of *PCK1* and *FOS* mRNAs, two established endogenous β2-AR transcriptional targets.^17,18^ Similar to the trends observed with the CREB reporter, transcriptional induction of *PCK1* and *FOS* mRNAs was significantly higher in the knockouts (Fig. 2c-d). Importantly, the results were also reproduced in cells, in which *RBM12* mRNA was depleted acutely by RNAi or CRISPR interference (CRISPRi)^19^ (Fig. S2a-d). Further, the trends were not selective for the full agonist isoproterenol, as we observed transcriptional hyperactivity after stimulation of knockout cells with a panel of endogenous and synthetic partial and full β2-AR agonists (Fig. 2e). Based on these data, both receptor-dependent PKA activity and transcriptional signaling are hyperactive in the absence of functional RBM12.

**Figure 2.**
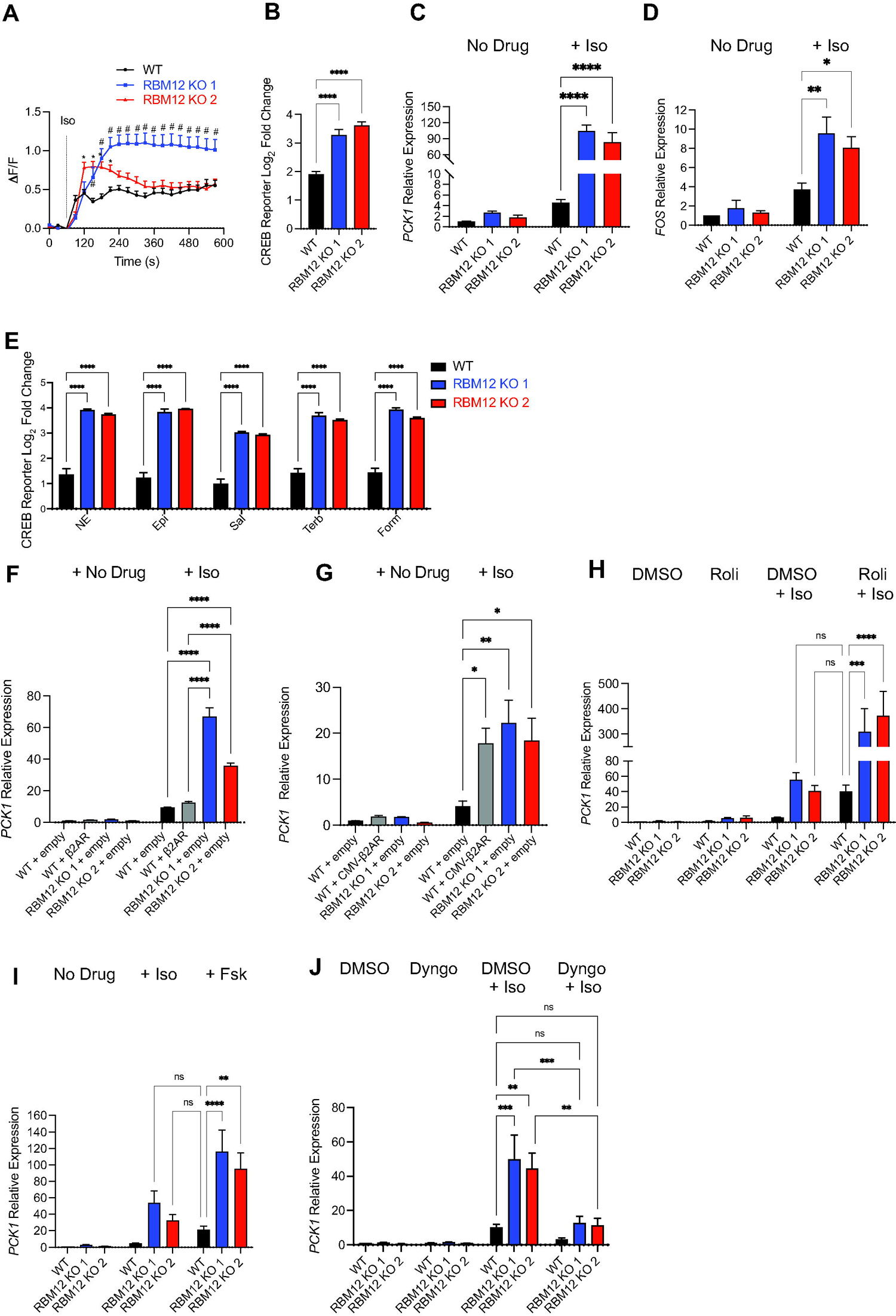
RBM12 loss leads to increased PKA activity and supraphysiological CREB-dependent transcriptional responses. (A) PKA sensor (ExRai-AKAR2) activity in response to 10 nM Iso (n = 24-47 cells from 3-4 independent transfections per cell line). # and * denote statistically significant timepoints between RBM12 KO1 vs. wild-type (WT) and KO2 vs. wildtype (WT), respectively. (B) Flow cytometry measurement of fluorescent CREB transcriptional reporter (CRE-DD-zsGreen) in response to 1 μM Iso and 1 μM Shield, 4 hours (n = 17). (C) RT-qPCR analysis of the endogenous β2-AR transcriptional target mRNAs, *PCK1* (n = 15) and (D) *FOS* (n = 9) in untreated cells or in cells treated with 1 μM Iso for 1 hour. (E) Flow cytometry measurement of the fluorescent CREB transcriptional reporter (CRE-DD-zsGreen) in response to a panel of endogenous (10 μM norepinephrine/NE, 10 μM epinephrine/Epi) or synthetic (10 μM salbutamol/Sal, 10 μM terbutaline/Terb, 50 nM formoterol/Form) β2-AR agonists and 1 μM Shield for 4 hours (n = 5). (F-G) *PCK1* mRNA expression in untreated or 1 μM Iso-treated cells transfected with empty plasmid (n = 3), (F) plasmid construct expressing lower levels of β2AR (n = 3), or (G) plasmid expressing β2AR from a CMV promoter (n = 3). (H) *PCK1* mRNA expression in untreated or 1 μM Iso-treated cells for 1 hour in the presence of either vehicle (DMSO) or 10 μM of the PDE4 inhibitor Rolipram (n = 4-5). (I) RT-qPCR of *PCK1* mRNA in untreated cells or cells treated with 1 μM Iso-treated cells or 10 μM forskolin for 1 hour (n = 12-13). (J) *PCK1* mRNA expression in cells pretreated with either vehicle (DMSO) or 30 μM Dyngo-4A for 20 min, then treated with 1 μM Iso for 1 hour (n = 5). All data are mean ± SEM. Statistical significance was determined using multiple unpaired *t*-tests with Benjamini, Krieger and Yekutieli false discovery rate correction (A), one-way ANOVA with Dunnett’s correction (B) or two-way ANOVA with Tukey’s correction (C-J). See also Figure S2.

We noted that the extent of β2-AR-dependent transcriptional hyperactivation observed in RBM12 knockout cells is striking, especially in the case of *PCK1* mRNA, which we have previously found to be the most robust transcriptional target of the receptor (~36-51-fold, Fig. 2c).^17^ We addressed whether this level of hyperactivation could be recapitulated in wild-type cells and, if so, under what biological conditions. First, we sought to overexpress the β2-AR reasoning that this would lead to increased cAMP production and transcriptional signaling. We began by expressing the receptor using a weak promoter. Specifically, we amplified a genomic region that included the ~1 kb sequence upstream of the *ADRB2* ORF, which contains the annotated 5’ UTR and promoter, and the gene coding sequence. We transiently transfected wild-type cells with the construct and observed that *ADRB2* mRNA levels were notably increased relative to endogenous *ADRB2* in empty vector-transfected cells (~25-fold increase) (Fig. S2e). This resulted in ~1.3-fold higher isoproterenol-dependent induction of the target mRNA *PCK1*. However, this increased transcriptional response did not match the striking effect seen upon RBM12 depletion (Fig. 2f). Next, we expressed the receptor under a CMV promoter leading to dramatically higher *ADRB2* mRNA levels relative to vector-transfected cells (~1000-fold) (Fig. S2f). Under these conditions, β2-AR-dependent *PCK1* mRNA upregulation resembled the levels seen in knockout cells (Fig. 2g). Based on these results, we hypothesized that loss of RBM12 may mimic a signaling state that can be induced only under supraphysiological activation conditions. To test whether this is indeed the case, we stimulated supraphysiological cAMP in wild-type cells with the following treatments: 1) saturating isoproterenol in the presence of the phosphodiesterase 4 (PDE4) inhibitor compound rolipram,^20^ or 2) direct activation of adenylyl cyclase with high doses of forskolin (10 μM). Both activation conditions yielded transcriptional signaling in wild-type cells that matched the levels observed in the knockouts (Fig. 2h-i). Therefore, RBM12 depletion leads to supraphysiological transcriptional signaling downstream of GPCR activation.

β2-AR signaling was recently found to be compartmentalized, with the β2-AR generating a second wave of signaling from endosomal membranes to selectively induce certain downstream responses. Specifically, the entire repertoire of transcriptional responses was induced by endosomal β2-AR signaling, while plasma membrane receptor activity was effectively uncoupled from this process.^17,21,22^ Given that loss of RBM12 results in higher cAMP and supraphysiological transcriptional responses, we next asked whether the compartmentalization of the GPCR signal may be impaired in the knockouts. To test if plasma membrane-resident β2-ARs may be capable of transducing transcriptional responses upon depletion of RBM12, we acutely blocked receptor internalization into endosomes with the dynamin inhibitor, Dyngo-4a.^23^ Consistent with previous reports, pretreatment with Dyngo-4a severely abolished *PCK1* mRNA induction by isoproterenol in wild-type cells (Fig. 2j).^17,21,22^ In marked contrast, β2-ARs confined to the cell surface of RBM12 knockout cells stimulated transcriptional responses that were on par with what was observed in wild-type cells with normal β2-AR trafficking (log_2_FC=3.21 ± 0.32 in vehicle-treated wild-type cells vs log_2_FC=2.94 ± 0.50 and 3.17 ± 0.57 in Dyngo-4a-treated knockout cell lines, respectively) (Fig. 2j, compare the “WT + Iso/DMSO to “KO + Iso/Dyngo” bars). These responses were further increased in vehicle-treated knockout cells stimulated with isoproterenol (Fig. 2j). Therefore, while transcriptional responses are orchestrated from endosomal β2-ARs in wild-type cells, both plasma membrane and endosomal β2-ARs stimulate gene expression when RBM12 is depleted. Importantly, this is not due to differences in receptor trafficking between the cell lines (Fig. S2g-h).

### The cAMP and transcriptional signaling steps are subject to discrete RBM12-dependent regulation

While we observed that loss of RBM12 impacted several steps in the GPCR cascade, it remained to be determined whether hyperactive cAMP production could account for the differences in the downstream responses, or whether RBM12 function impacts multiple steps in the pathway.

To address this, we first sought to carefully match isoproterenol activation conditions to yield comparable induced cAMP levels between wild-type and knockout cells, and then examined the resulting downstream transcriptional activation. We reasoned that this would allow us to separate the impact of RBM12 depletion on cAMP production from that on downstream transcriptional signaling. Stimulation of knockouts with sub-saturating doses of isoproterenol (10 nM) led to cAMP accumulation that was equivalent to that generated by stimulation of the wild-type with saturating isoproterenol (Fig. S3). Yet, these matched cAMP stimulation conditions still yielded significantly higher transcriptional signaling in RBM12-depleted cells (6.3-9-fold higher *PCK1* mRNA induction compared to wild-type, *p*-value < 1.00 x 10^-4^ by oneway ANOVA test) (Fig. 3a). To complement these experiments, we next bypassed adenylyl cyclase activation altogether by treating cells with a permeable cAMP analog, 8-CPT-cAMP.^24^ We observed hyperactive transcriptional signaling in knockout cells reflected by increased 8-CPT-cAMP-dependent *PCK1* mRNA induction (Fig. 3b). Collectively, these results suggest that cAMP production and transcriptional signaling are independently subject to RBM12 regulation.

**Figure 3.**
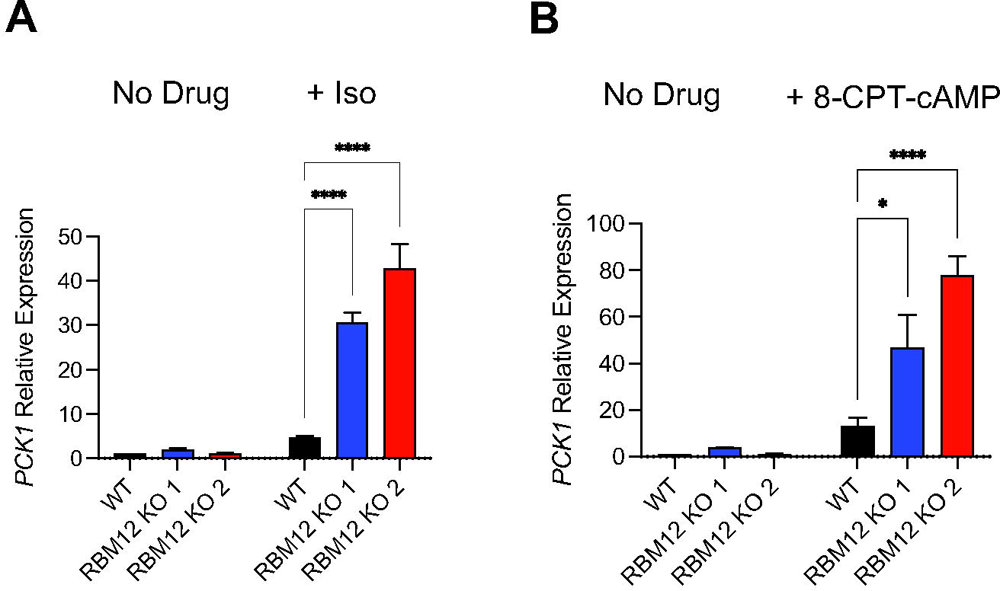
RBM12 independently impacts the cAMP and transcriptional signaling steps. (A) *PCK1* mRNA expression in untreated cells or in cells treated with either 1 μM isoproterenol (wild-type cells) or 10 nM isoproterenol (RBM12 knockout cells) for 1 hour (n = 3). (B) *PCK1* mRNA expression in untreated cells or in cells treated with 150 μM 8-CPT-cAMP for 1 hour (n = 4). All data are mean ± SEM. Statistical significance was determined using two-way ANOVA with Tukey’s correction (A-B). See also Figure S3.

### The neuropsychiatric disease-linked mutations fail to rescue GPCR-dependent hyperactivation in cells depleted of RBM12

Two truncating variants in the *RBM12* gene (c.2377G>T and c.2532delT) have been implicated as genetic risk factors for familial psychosis.^11^ However, it is not known how either mutation impacts protein function. We speculated that these mutations may result in nonfunctional RBM12 and therefore impact GPCR/cAMP signaling responses. To test this, we asked whether constructs encoding either wild-type or variant RBM12 could genetically rescue the hyperactive transcriptional signaling in RBM12-depleted cells. We chose to deplete *RBM12* expression by CRISPRi. This approach utilizes a catalytically dead Cas9 fused to a transcriptional repressor to target endogenous promoter regions and therefore enables rescue with constructs driven by artificial promoters.^13^ To express RBM12 variants, we cloned lentiviral plasmids encoding N-terminally EGFP-tagged wild-type (WT), c.2377G>T (‘G>T’), or c.2532delT (‘delT’) RBM12 expressed under the mammalian Ubiquitin C promoter. First, we established that these constructs generated proteins of expected sizes by Western blot. We observed that the two truncating mutations gave rise to smaller protein products. In addition, we noted that the delT variant resulted in reduced protein expression compared to WT and G>T (Fig. 4a, compare lanes 1 and 2 to lane 3). We did not observe downregulation in delT mRNA levels (Fig. S4a), suggesting that the mutation impacts a post-transcriptional step, either protein translation or stability. Next, we tested whether the mutant proteins displayed altered subcellular localization. Studies done in primary human erythroblasts, A-431, and U-2OS cells found that RBM12 displayed nuclear localization.^9,25^ In agreement with these reports, we observed nuclear localization for native RBM12 under both basal and isoproterenol-induced conditions in HEK293 cells (Fig. 4b). Similarly, wild-type EGFP-RBM12 localized to the nucleus (Fig. 4c, top panel). The truncating mutations did not perturb protein localization as both the G>T and delT variants also concentrated in the nucleus (Fig. 4c, middle and bottom panels).

**Figure 4.**
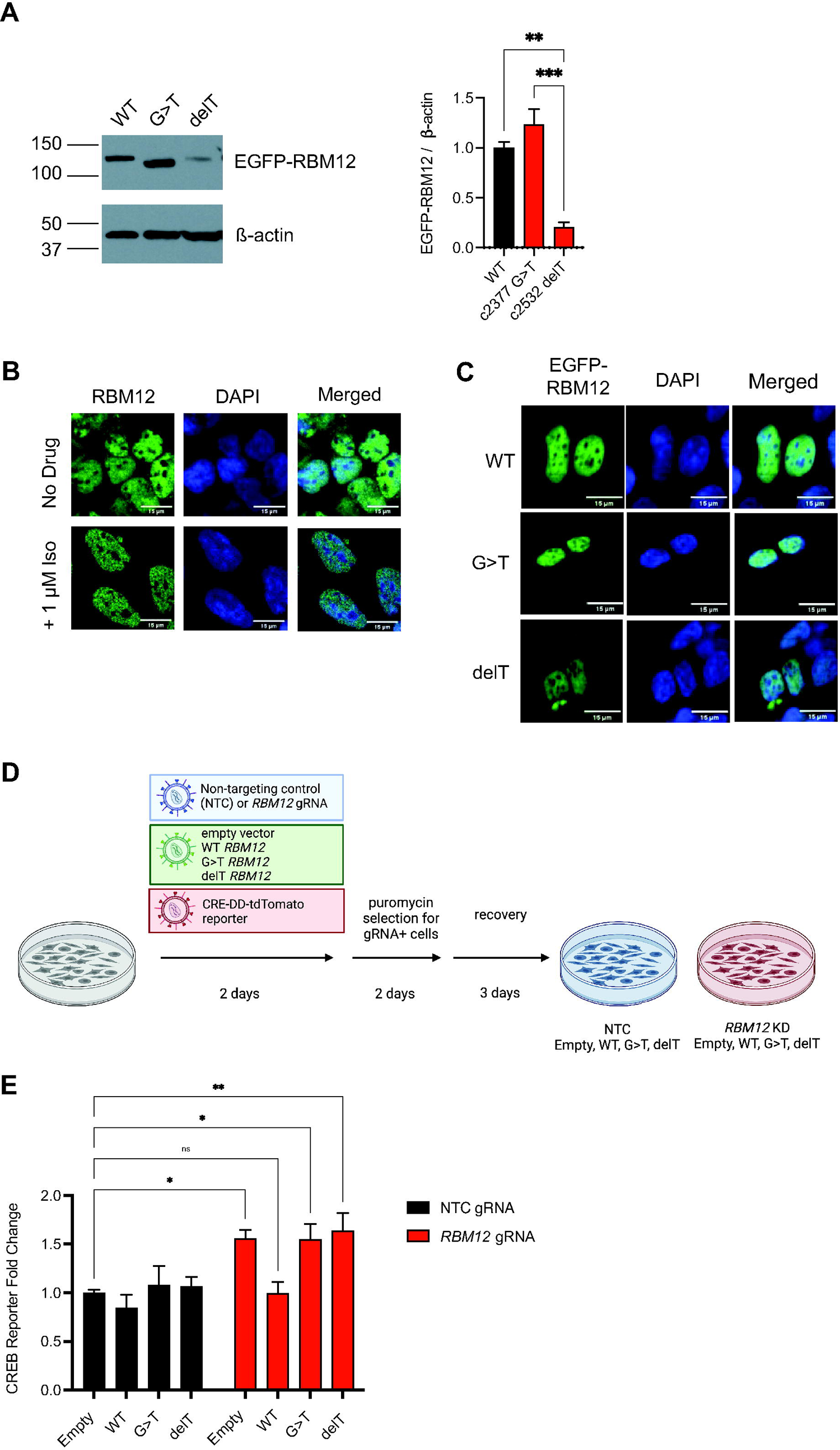
Expression of disease-associated variants in RBM12 knockdown cells does not rescue the hyperactive GPCR-dependent transcriptional signaling. (A) Western blot analysis of EGFP-tagged wild-type, c.2377G>T and c.2532delT RBM12 (n = 3) probed with antibody recognizing EGFP. All data are normalized relative to wild-type. (B) Fixed cell fluorescence microscopy analysis of endogenous RBM12 in untreated and stimulated (1 μM Iso for 20 minutes) HEK293 cells. (C) Localization of EGFP-tagged wild-type, c.2377G>T or c.2532delT RBM12 by fluorescence microscopy. (D) Schematic of the flow cytometry-based rescue experiment. (E) Flow cytometry measurement of the fluorescent CREB transcriptional reporter (CRE-DD-tdTomato) in response to 1 μM Iso and 1 μM Shield for 6 hours (n = 8-11). Data are normalized relative to the “NTC + empty vector” sample values. All data are mean ± SEM. Statistical significance was determined using one-way ANOVA (A) or two-way ANOVA with Dunnett’s correction (E). See also Figure S4.

We proceeded to evaluate whether the mutants could rescue the hyperactive GPCR/cAMP signaling phenotype seen in RBM12-depleted cells. We simultaneously transduced HEK293 cells stably expressing dCas9-KRAB with 1) either a non-targeting control (NTC) gRNA or a gRNA against *RBM12*, and 2) an empty EGFP backbone, WT, G>T, or delT *RBM12* constructs. To monitor CREB-dependent transcription, we generated a new version of the CREB transcriptional reporter, in which cAMP responsive elements drive the production of a red fluorescent protein in response to cAMP accumulation. This allowed us to selectively examine reporter levels in cells that express the CRISPRi gRNA (tagged with BFP) and the RBM12 constructs (tagged with EGFP) by flow cytometry (Fig. 4d and Fig. S4b-c). As expected, we observed increased reporter accumulation upon isoproterenol stimulation in *RBM12* gRNA-transduced compared to NTC-transduced cells (Fig. 4e, leftmost black vs red bars). Notably, we found that normal signaling downstream of β2-AR activation was restored only in cells expressing WT, but not the G>T or delT mutant (Fig. 4e). Therefore, neither of the two disease-linked RBM12 variants could rescue impaired GPCR-dependent signaling.

### RBM12 is a repressor of GPCR/cAMP signaling in human iPSC-derived neurons

RBM12 is expressed ubiquitously across different tissues.^26^ However, it likely has important functions in the brain based on its ties to neuropsychiatric disorders^11^ and neurodevelopment.^12^ Therefore, to examine the role of RBM12 in GPCR/cAMP signaling in a physiologically relevant model, we chose human induced pluripotent stem cell-derived neurons (iNeurons). iNeurons offer a unique setting to investigate receptor activity in live human neurons, and a suitable genetic context to study the functional effects of disease-relevant gene mutations. Since glutamatergic signaling is of key significance in the pathobiology of neuropsychiatric diseases, including schizophrenia,^27–31^ we selected a previously published clonal hiPSC line harboring an inducible Neurogenin 2 *(NGN2)* driven by a doxycycline-inducible promoter for glutamatergic differentiation.^32^ In addition, this line expresses dCas9-Krab-BFP and is therefore compatible with CRISPRi-based gene silencing (Fig. 5a). We observed nuclear staining of native RBM12 under both basal and isoproterenol-induced conditions in neurons, consistent with its localization in HEK293 cells (Fig. 5b). Next, to deplete *RBM12*, we utilized either NTC or *RBM12*-targeting gRNAs, sorted the gRNA-transduced cells, and confirmed *RBM12* mRNA depletion by qPCR and protein expression by Western blotting (Fig. S5a). Importantly, we verified that these cells expressed comparable levels of FLAG-β2-AR upon differentiation into iNeurons (Fig. S5b).

**Figure 5.**
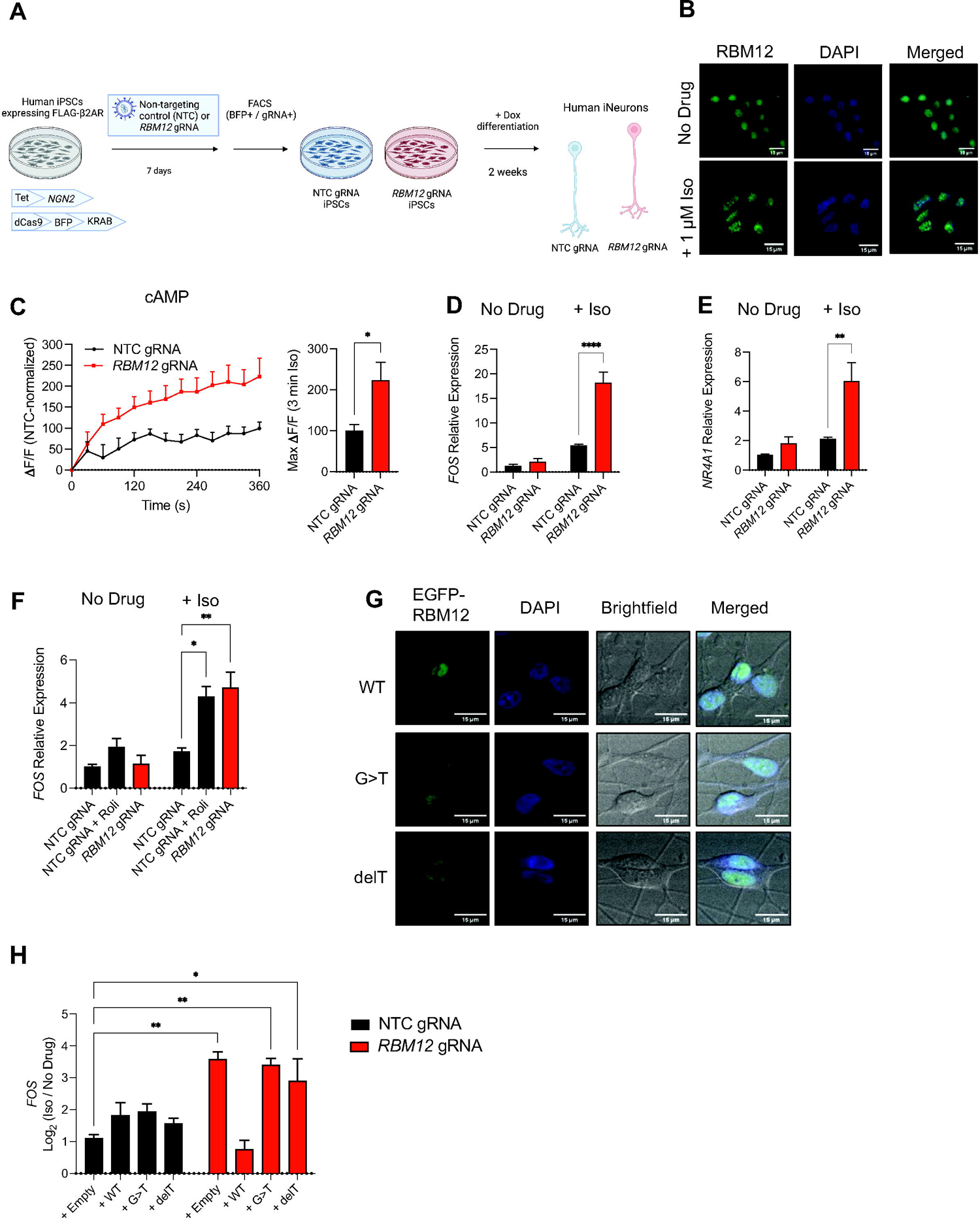
Signaling hyperactivity upon loss of RBM12 in human iPSC-derived neurons. (A) Schematic of the CRISPRi-mediated RBM12 depletion in human iPSC-derived neurons. (B) Fixed cell fluorescence microscopy imaging of native RBM12 in untreated and stimulated (1 μM Iso for 30 minutes) iNeurons. (C) Accumulation of the fluorescent cADDis sensor in NTC- and *RBM12* gRNA-expressing neurons in response to 1 nM Iso (n = 6). (D) Expression of *FOS* and (E) *NR4A1* mRNAs in response to treatment with 1 μM Iso for 1 hour by RT-qPCR (n = 6). (F) Expression of *FOS* mRNA in response to 1 μM Iso in the presence of either vehicle (DMSO) or 10 μM Rolipram in neurons treated for 1 hour (n = 3). (G) Expression of EGFP-tagged wild-type, c.2377G>T or c.2532delT RBM12 by fluorescence microscopy. (H) Expression of *FOS* mRNA in response to stimulation with 1 μM Iso for 1 hour in neurons transduced with empty plasmid or plasmid encoding WT, G>T, or delT EGFP-RBM12 (n = 2-3). All data are mean ± SEM. Statistical significance was determined using unpaired t-test (C) or two-way ANOVA with Tukey’s correction (D-F, H). See also Figure S5.

We began by differentiating hiPSCs with doxycycline for 2 weeks and examining the impact of *RBM12* depletion on cAMP accumulation. We used an established fluorescent cytosolic cAMP biosensor, cADDis.^33^ We found that iNeurons transduced with *RBM12* gRNA exhibited hyperactive cAMP signaling upon β2-AR activation relative to NTC-harboring iNeurons (Fig. 5c). To examine transcriptional responses, we utilized RT-qPCR analysis of *NR4A1* and *FOS* mRNAs, two known CREB-dependent immediate early genes induced by neuronal activity.^34–37^ We saw that *RBM12* knockdown yielded hyperactive upregulation of both mRNAs (Fig. 5d-e). Notably, the extent of transcriptional hyperactivity seen in the knockdown could be recapitulated in NTC-harboring cells under supraphysiological cAMP activation with isoproterenol and rolipram (Fig. 5f). Lastly, we assessed whether the two RBM12 mutants could rescue the impaired signaling phenotype. Wild-type (WT), G>T, and delT RBM12 transduced into iNeurons localized to the nucleus (Fig. 5g). In agreement with the expression trends seen in HEK293, we observed reduced delT mutant expression compared to WT or G>T RBM12 (Fig. S5c-d). RBM12-dependent hyperactive transcriptional signaling was abolished upon expression of WT RBM12, but not G>T or delT, as evaluated by RT-qPCR analysis of *FOS* mRNA (Fig. 5h). Thus, depletion of *RBM12* in human neurons recapitulates the gamut of signaling phenotypes seen in HEK293. Collectively, these results support conserved RBM12-dependent regulation of the GPCR/cAMP pathway across cell types, including in a physiologically relevant system.

### β2-AR activation in RBM12-depleted cells leads to modified neuronal transcriptional responses

To capture the cellular functions impacted by *RBM12* depletion, we carried out genome-wide transcriptomic analysis of untreated and isoproterenol-activated NTC- and *RBM12* gRNA-expressing iNeurons. We began by defining a set of all possible β2-AR-dependent transcriptional targets across the two neuronal lines. For this analysis, we independently identified target sets in NTC-transduced (“wild-type”) and *RBM12* knockdown neurons by differential expression analysis between each respective basal and isoproterenol conditions. We obtained a total of 669 unique β2-AR-dependent transcriptional targets across the two cell lines (Fig. 6a, Table 1). To elucidate specific β2-AR-dependent neuronal processes, we performed Gene Ontology (GO) Term analysis of the targets.^38^ We identified a breadth of regulated processes including metabolic processes, cell differentiation, response to hormone/stimulus, regulation of gene expression, learning and memory, and neuronal differentiation (FDR *p*_adj_ < 5.0×10^-2^ by Fisher’s exact test) (Fig. 6b, Table 2). Importantly, at least 110 genes associated with neuronal activity, memory, cognition, as well as synaptic transmission and synapse formation were represented among the β2-AR-dependent targets.^37^ These included brain-derived neurotrophic factor (*BDNF*) and transcription factor-coding genes (e.g., *NR4A1, FOS, FOSB, EGR1, JUN*), consistent with the reported critical roles of the β2-AR/cAMP pathway in neurobiology (Table 1).^39^

**Figure 6.**
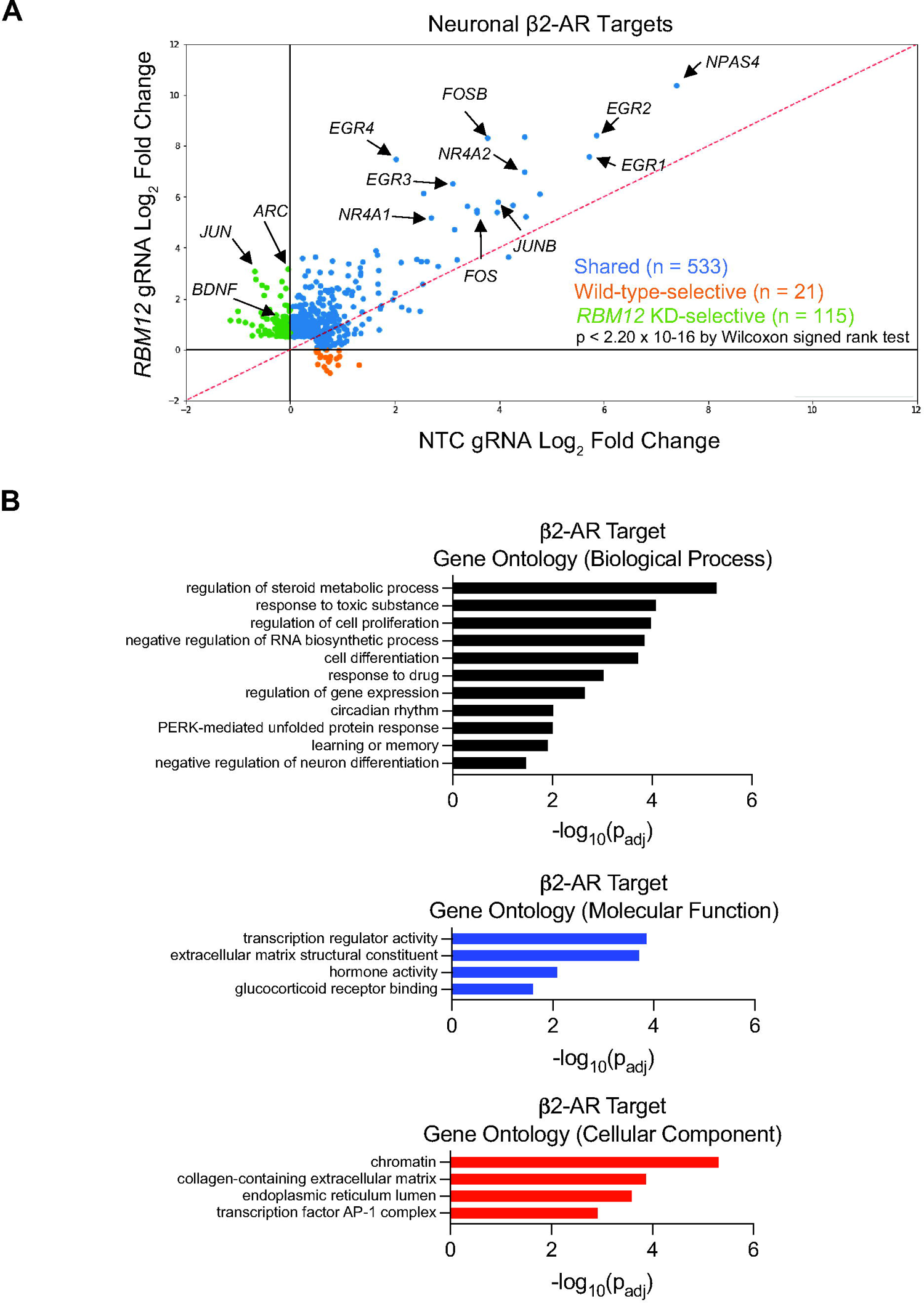
RBM12 loss impacts the β2-AR neuronal transcriptional responses. (A) Scatter plot showing gene fold induction (log_2_ Iso/No Drug) of neuronal β2-AR targets (n = 669) identified by RNA-seq analysis (n = 3 per cell line per drug condition). Blue dots represent genes that were induced by a 1-hour treatment with 1 μM Iso in both wild-type (NTC gRNA) and *RBM12* KD *(RBM12* gRNA) neurons. Orange dots represent genes that were induced only in wild-type and unchanged or downregulated in *RBM12* KD neurons. Green dots represent genes that were induced only in *RBM12* KD neurons and unchanged or downregulated in wild-type. Indicated by arrows are a subset of genes with established roles in neuronal activity. The underlying information is summarized in Table 1. (B) Gene Ontology categories enriched among the neuronal β2-AR targets from (A). The underlying information is presented in Table 2. See also Figure S6.

Under basal conditions, we did not observe any notable trends toward RBM12-dependent increase in the expression of the targets (Fig. S6a). However, under isoproterenol stimulation, we found significant transcriptional hyperactivation across all target genes in the *RBM12* knockdown (*p* < 2.20 x 10^-16^ by Wilcoxon signed rank test). To gain deeper insight into these differences, we next asked how many of the targets are selectively induced in response to β2-AR stimulation in each cell line. In principle, genes may be upregulated in either wild-type or *RBM12* knockdown, representing qualitatively distinct targets. On the other hand, targets may be changing in both wild-type and *RBM12*-depleted neurons but to different extent (quantitatively distinct targets). To identify qualitatively unique targets for each cell line within the set of 669 genes, we applied a cut-off of gene log_2_-fold change ≥ 0 with isoproterenol in one neuronal line but not the other. This analysis yielded 21 wild-type- and 115 *RBM12* knockdown-specific target genes (Table 1). Interestingly, genes with known functions in synapses, memory and cognition were represented in both exclusive lists. Factors involved in synaptic plasticity such as *JUN, ARC* (encoding the activity-regulated cytoskeleton-associated protein), *BDNF*, and *NRXN3* (encoding the cell adhesion molecule neurexin-3-alpha) were induced only in *RBM12* knockdown neurons, while *GRIA2* (encoding the AMPA receptor) and *CBLN2* (encoding cerebellin 2 precursor) were upregulated only in NTC neurons (Fig. 6a, Table 1). While the remaining 533 genes were induced in both cell lines, there was a significant trend toward RBM12-dependent hyperactivation (*p* < 2.20 x 10^-16^ by Wilcoxon signed rank test). These results indicate that the neuronal β2-AR-dependent transcriptome is disrupted by *RBM12* depletion, resulting in both quantitatively and qualitatively distinct responses.

### RBM12 impacts GPCR/cAMP signaling through regulation of adenylyl cyclase and PKA expression

RBM12 is localized in the nucleus (Fig. 4b) and was recently shown to have RNA-binding activity.^4^ Yet, loss of RBM12 impacts multiple signaling steps, including cAMP production which takes place in the cytosol. Thus, we speculated that the observed regulation may take place through other factor(s), that in turn are dependent on RBM12 for proper expression or function. To begin to dissect the mechanisms governing the RBM12-dependent regulation of GPCR signaling, we performed differential expression analysis between untreated wild-type and untreated *RBM12*-depleted neurons. We identified 2,645 differentially expressed genes (Table 3). Gene Ontology analysis identified processes related to nervous system development and function as well as “GPCR signaling pathway” and “adenylyl cyclase - modulating signaling pathway” (Fig. 7a, Table 3). We were particularly intrigued by the latter GO term categories, as they could provide insights into the identity of the proximal regulator(s) of our pathway. We found altered expression of several effectors with known roles in the GPCR/cAMP cascade, including adenylyl cyclases, PDEs and PKA isoforms (Table 3). We sought to further narrow down the factors by identifying effectors that display consistent RBM12-dependent abundance changes in both iNeurons and HEK293 cells. For that, we carried out transcriptomic analysis of the HEK293 knockout and parental lines and compared the list of differentially expressed genes between model systems to find shared cAMP pathway effectors. For genes expressed in both models, we required that 1) factor abundance changes trended in agreement in the neuron and HEK293 experiments, and 2) these changes were statistically significant (p_adj_ < 5.0×10^-2^ by Wald test) in at least 2 out of the 3 depleted cell line models. As a result, we found that two adenylyl cyclase isoforms *(ADCY3* and *ADCY5)* were consistently upregulated, while *PDE7A* was downregulated (Fig. 7b). In addition, the abundance of the neuron-specific *ADCY8* and *PDE1C* was aberrant: *ADCY8* levels were increased, while *PDE1C* levels were decreased upon *RBM12* depletion in neurons (Fig. 7b). The fact that *ADCY* abundance was higher in *RBM12*-depleted cells suggests that, in addition to stimulated cAMP, basal cAMP levels may also be increased. Indeed, we found that cAMP concentrations were significantly elevated in untreated RBM12 knockout relative to wild-type HEK293 cells (Fig. 7c).

**Figure 7.**
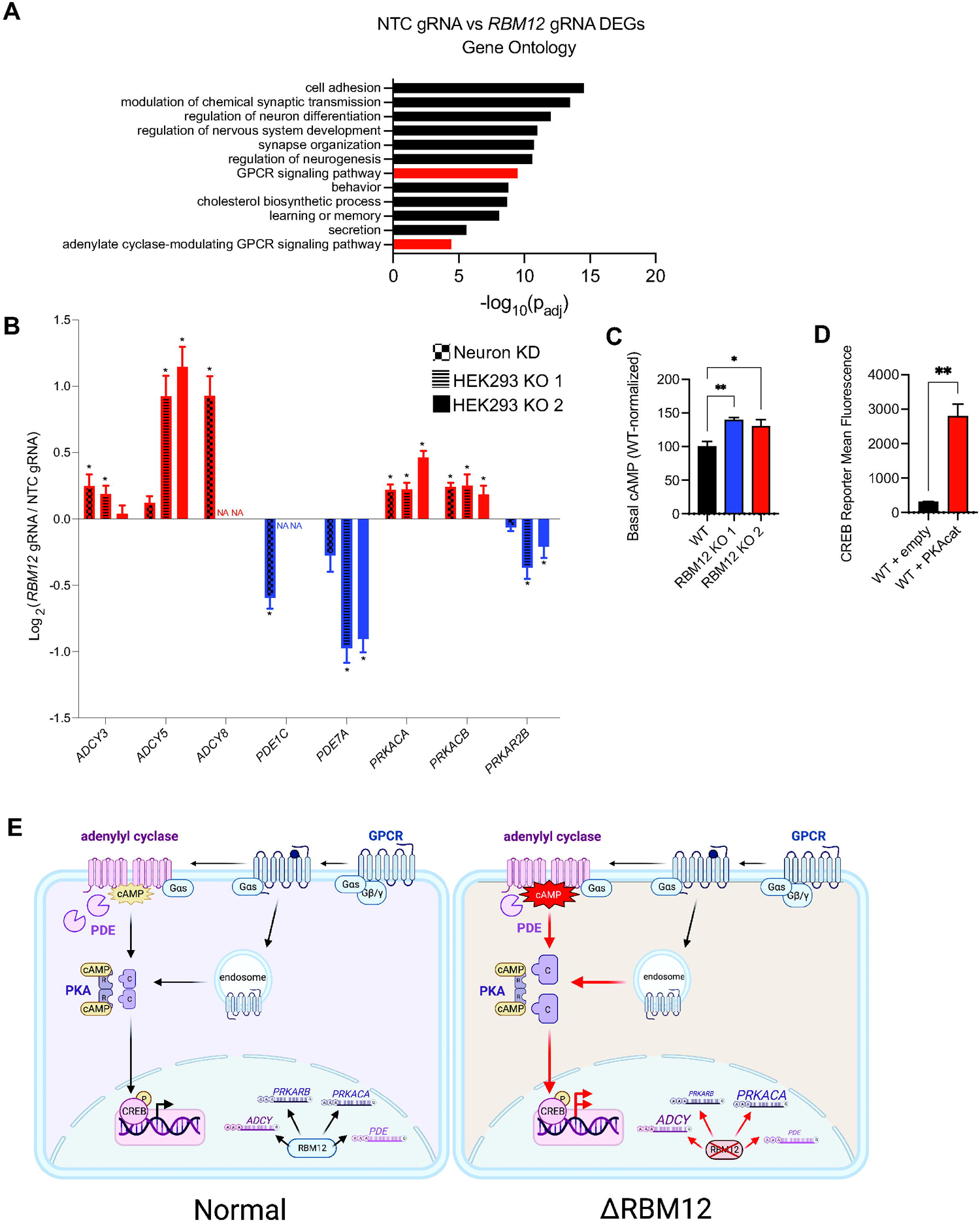
RBM12 regulates the expression of multiple GPCR/cAMP effectors. (A) Gene ontology analysis of differentially expressed genes between wild-type and *RBM12* KD neurons (n = 2,645 genes). (B) Graph summarizing abundance changes (log_2_ fold change RBM12 gRNA/NTC gRNA) of known GPCR/cAMP regulators in wild-type and *RBM12* KD neurons (n = 3 per cell line) and HEK293 (n = 7 in wild-type cells and n = 3 per knockout cell line). Asterisks denote statistical significance (*p*_adj_ < 5.0 x 10^-2^ by Wald test). “NA” denotes genes not expressed in HEK293. (C) Basal cAMP levels in wild-type and HEK293 RBM12 knockouts measured using ELISA assay (n = 5). All values are normalized relative to wild-type. (D) Flow cytometry measurement of the fluorescent CREB transcriptional reporter (CRE-DD-zsGreen) in wild-type cells expressing either empty plasmid or plasmid encoding PKAcat following stimulation with 1 μM Iso and 1 μM Shield for 4 hours (n = 3). (E) Model of RBM12-dependent regulation of the GPCR/cAMP signaling pathway. All data are mean ± SEM. Statistical significance was determined using adjusted *p*-value corrected for multiple testing by Wald test (B), one-way ANOVA with Dunnett’s correction (C), or unpaired t-test (D).

Furthermore, the expression of *PRKACA* and *PRKACB* which encode the PKA catalytic (PKAcat) subunits Cα and Cβ, were upregulated in RBM12-depleted cells (Fig. 7b). On the other hand, the expression of *PRKAR2B*, which encodes the PKA type II-beta regulatory subunit (RIIβ), was downregulated in RBM12-depleted cells (Fig. 7b). Mirroring the trends in mRNA, the protein levels of the PKAcat isoform predominantly expressed in most tissues,^40,41^ Cα, were also higher in both RBM12 knockout HEK293 cells and in neurons acutely depleted of *RBM12* by CRISPRi (Fig. S6b-c). To test whether PKA over-expression alone is sufficient to induce hyperactive downstream transcription, we transiently expressed PKAcat in wild-type HEK293 cells and measured fluorescent CREB reporter levels. We found that β2-AR-dependent accumulation of the reporter was indeed higher in cells transfected with PKAcat (Fig. 7d). Therefore, the RBM12-dependent transcriptional hyperactivity, including following cascade stimulation with a cAMP analog (Fig. 3b), could be driven at least in part by increased expression of PKAcat. Collectively, the transcriptomic changes induced by loss of RBM12 are consistent with the hyperactive GPCR/cAMP/PKA signaling phenotypes, and support impaired *ADCY*, *PDE*, and *PRKACA* abundance as an underlying molecular mechanism.

## Discussion

In this study, we identify and characterize RBM12 as a novel repressor of GPCR/cAMP signaling. Although mutations in *RBM12* have been linked to heritable psychosis and neurodevelopmental defects,^11,12^ its precise cellular functions are not well-understood. We discover that RBM12 independently impacts multiple steps within the GPCR pathway. Loss of RBM12 leads to hyperactive basal and stimulated cAMP (Fig. 7c and Fig. 1c) as well as increased PKA activity and transcriptional signaling upon activation of the β2-AR (Fig. 2).

RBM12 is predominantly localized to the nucleus^25^ and has been shown to bind RNA and to tentatively impact transcript splicing.^9^ Based on the present work, we cannot rule out that the cytosolic fraction of RBM12 plays a direct regulatory role in GPCR signaling. Studies aimed at defining the RBM12 interactome could resolve this possibility. However, we favor a more parsimonious model, according to which nuclear RBM12 controls the transcriptional or post-transcriptional fate of proximal cytosolic factors that in turn directly impact the GPCR pathway (Fig. 7e). In support of this, we demonstrate that loss of RBM12 leads to impaired expression of multiple factors of established significance within the GPCR/cAMP cascade, including adenylyl cyclases, PDEs, and PKA regulatory and catalytic subunits (Fig. 7b). Upon RBM12 loss, PKA RIIβ levels are decreased, while Cα and Cβ levels are increased. This result is intriguing in the context of our current understanding of how PKAcat diffusion may be limited in cells. The PKA regulatory subunits are in large molar excess of the catalytic subunits in a wide range of tissues, including in the brain, facilitating efficient ‘recapture’ of liberated catalytic subunits.^42^ Based on this, we speculate that depletion of RBM12 likely leads to reduced anchoring of PKAcat and therefore to loss of compartmentalized PKA activity. Impaired compartmentalization of GPCR and cAMP signaling would be expected to arise also from the altered adenylyl cyclase and PDE abundance in *RBM12*-depleted cells. Several *ADCY* genes are upregulated, while *PDE* isoforms are downregulated in *RBM12* knockdowns (Fig. 7b). Indeed, we report that transcriptional signaling, a cellular response downstream of the β2-AR that is known to be spatially encoded,^17,21,22^ is compromised in the knockout (Fig. 2j). Furthermore, the hyperactive transcriptional response seen in the mutant can be recapitulated only under supraphysiological activation conditions, including PDE inhibition and receptor overexpression (Fig. 2h-i), which are expected to abolish compartmentalized signaling.^17,43,44^ Future studies that apply cAMP and PKA biosensors targeted to distinct subcellular organelles would be invaluable in further defining local signaling in RBM12 knockouts.

Two mutations in the *RBM12* gene (c.2377G>T and c.2532delT), each resulting in the truncation of the terminal RRM domain, were recently associated with familial psychosis.^11^ Here, we observe that the delT protein is expressed at decreased levels compared to wildtype or G>T RBM12 (Fig. 4a, S4b, S5d). We further find that neither variant can rescue aberrant GPCR/cAMP signaling when expressed in HEK293 cells and neurons depleted of RBM12 (Fig. 4e and 5h). Accordingly, the two mutations lead to loss-of-function protein likely through distinct mechanisms. While the delT-related phenotypes may stem at least in part from diminished protein expression, the G>T mutation may interfere with RBM12 function by abolishing interactor binding. Interestingly, the last RRM domain does not appear to be required for all processes regulated by RBM12. A recent study found a novel role of RBM12 as a repressor of fetal hemoglobin expression.^9^ Notably, the authors reported a requirement only for the N-terminal portion of the protein in that process, whereas deletion of the C-terminal RRM domain, which is truncated in the psychosis cohorts, was dispensable from that regulation. Therefore, the C-terminal RRM domain and its interactome must be of particular significance in the context of RBM12 and its role in GPCR/cAMP signaling. We note that our study is limited by the over-expression of RBM12 variants in a genetically depleted background, and thus may not fully recapitulate the consequences of the disease-relevant mutations. Future work using endogenous systems, including cells derived from patients harboring the mutations or CRISPR knock-in strategies to generate heterozygous mutant cell lines, would offer more direct and detailed insights into their functions.

It is tempting to speculate that dysregulation of GPCR signaling could be one important molecular pathway that contributes to the neuronal pathologies stemming from loss of RBM12. More than a third of all known GPCRs are expressed in the brain, where they bind to neurotransmitters and play essential roles in synaptic and structural plasticity.^45–47^ Dysregulation of GPCR activity in the brain also contributes to the pathophysiology of several neurological and neuropsychiatric disorders.^48–50^ Similarly, cAMP is a critical second messenger that mediates all important aspects of neuronal function, including development, excitability, and plasticity.^51^ While most of this work is centered around the prototypical β2-AR, we report that RBM12 function is required for normal cAMP production downstream of other Gαs-coupled receptors with established roles in the nervous system (dopamine 1 and adenosine receptors, Fig. 1d-e).^14,52^ This finding is perhaps not surprising given that all stimulatory GPCRs converge on the same components of the cAMP/PKA pathway, many of which are dysregulated in the absence of RBM12 (Fig. 7b). As a result, the neuronal GPCR-driven transcriptional responses are altered when *RBM12* is knocked down. Specifically, the entire repertoire of targets, many of which orchestrate processes essential for neuronal differentiation, gene reprogramming, and memory and learning, shows a trend towards hyperactivation in depleted neurons (Fig. 6a). Moreover, more than 100 genes are induced in response to receptor stimulation only in the knockdown. Most notable among this set are *ARC* and *BDNF*, two factors with crucial roles in synaptic function, plasticity, and learning (Table 1).^53,54^ At the same time, we cannot rule out the possibility that RBM12 impacts processes in the brain through additional mechanisms. In fact, we identify extensive reprogramming of RBM12 knockdown neurons beyond expression of cAMP pathway effectors, as the set of altered GO categories includes neuron differentiation, synapse organization, and neurogenesis (Fig. 7a, Table 3). It remains to be determined how all these changes collectively impact neuronal function, and to what extent any of the alterations are driven specifically by dysregulation of the receptor pathway. Further, a different *RBM12* mutation has been implicated in the aberrant development of the mouse forebrain and midbrain.^12^ Therefore, investigation of the neurodevelopmental consequences of RBM12 depletion would likewise be warranted.

The cAMP/PKA cascade has been implicated in neuropsychiatric disorders in the past supporting the requirement for tightly regulated cAMP signaling for proper neuronal function. A study on post-mortem brains of patients with bipolar affective disorder demonstrated elevated levels of the PKAcat subunit Cα in temporal and frontal cortices compared to matched normal brains.^55,56^ A different report on patient-derived platelet cells found that the catalytic subunit of cAMP-dependent protein kinase was significantly upregulated in untreated depressed and manic patients with bipolar disorder compared with untreated euthymic patients with bipolar disorder and healthy subjects.^57^ In the context of schizophrenia, cAMP/PKA signaling has been found to be both reduced and hyperactive. For example, the PKA regulatory subunits, RI and RII, were significantly reduced in platelets from schizophrenic patients in one report,^58^ and several adenylyl cyclase isoforms were downregulated in patient fibroblasts reprogrammed into iPSC neurons in another.^59^ While this manuscript was under preparation, a study reported similar findings to ours with respect to the tentative molecular pathways dysregulated in loss-of-function mutants of another risk factor associated with neurodevelopmental defects and schizophrenia. Specifically, the authors found that heterozygous mutations in the histone methyltransferase SET domain-containing protein 1 A (*SETD1A*) led to transcriptional and signaling signatures supporting hyperactivation of the cAMP pathway through upregulation of adenylyl cyclases and downregulation of PDEs.^60^ This in turn resulted in increased dendritic branching and length and altered network activity in human iPSC-derived glutamatergic neurons.^60^ Therefore, the cAMP/PKA pathway appears to be a common point of convergence downstream of different risk factors for neuropsychiatric disorders and could present a therapeutic target in certain genetic contexts.

In summary, this study identifies a previously unappreciated role for RBM12 in the context of the GPCR/PKA/cAMP pathway. Because the regulation is conserved across multiple receptors and in two different cell models, it is likely of broad relevance and should be explored further as a tentative molecular mechanism underlying the functions of RBM12 in brain physiology and pathophysiology.

## Acknowledgments

We thank members of the Tsvetanova lab for valuable discussion and critical feedback on the manuscript. We thank the Duke Cancer Institute Flow Cytometry Shared Resource and the Duke University Light Microscopy Core Facility. We thank Jin Zhang (University of California, San Diego, CA) for providing the TagBFP-PKAcat and ExRai-AKAR2 constructs. We thank Mark von Zastrow (University of California, San Francisco, CA) for providing the pcDNA 3.1-D2R, pcDNA 3.1-Δ- and μ-opioid receptors, and FHUGW-EGFP constructs. Figures 4d, 5a and 7e were created using BioRender. This work was supported by the National Institutes of Health (R35GM142640 to N.G.T.), the American Association of University Women (2021 International Fellowship to K.M.S.), and the American Heart Association (Pre-Doctoral Fellowship #898739 to K.M.S.).

## Author Contributions

Conceptualization, N.G.T., K.M.S.; methodology, N.G.T., K.M.S., A.G.; investigation, N.G.T., K.M.S., and A.G.; formal analysis, N.G.T., K.M.S., and A.G.; funding acquisition, N.G.T. and K.M.S.; supervision, N.G.T.; writing – original draft, N.G.T., K.M.S.; writing – review & editing, N.G.T., K.M.S., and A.G.

## Table Legends

**Table 1.** Neuronal β2-AR transcriptional target genes identified by RNA-seq (669 genes).

**Table 2.** Gene Ontology analysis of neuronal β2-AR transcriptional targets.

**Table 3.** Differential expression of wild-type and *RBM12* knockdown neurons and Gene Ontology analysis.

## Materials and Methods

### Chemicals

(-)-Isoproterenol hydrochloride was purchased from Sigma-Aldrich (Cat. #I6504), dissolved in 100 mM ascorbic acid to 10 mM stock, and used at indicated concentrations. Alprenolol hydrochloride was purchased from Sigma-Aldrich (Cat. #A0360000), dissolved in DMSO to 10 mM stock, and used at 10 μM final concentration. Norepinephrine was purchased from Sigma (Cat. #A7257), dissolved in 100 mM ascorbic acid to 10 mM stock, and used at 10 μM final concentration. (-) Epinephrine was purchased from Sigma (Cat. #E4250), dissolved in 100 mM ascorbic acid to 10 mM stock, and used at 10 μM final concentration. Salbutamol was purchased from Cayman Chemical Company (Cat. #21003), dissolved in DMSO to 10 mM stock, and used at 10 μM final concentration. Formoterol fumarate dihydrate was purchased from Sigma Aldrich (Cat. #F9552), dissolved in DMSO to 10 mM stock, and used at 50 nM final concentration. Terbutaline hemisulfate salt was purchased from Sigma-Aldrich (Cat. #T2528), dissolved in water to 10 mM stock, and used at 10 μM final concentration. 5’-(N-Ethyl Carboxamide) adenosine (NECA) was purchased from Sigma-Aldrich (Cat. #119140), dissolved in DMSO to 10 mM stock, and used at 10 μM final concentration. SKF-1297 hydrobromide was purchased from Tocris (Cat. #1447), dissolved in DMSO to 10 mM stock, and used at 10 nM final concentration. ICI-118,551 hydrochloride was purchased from Sigma-Aldrich (Cat. #I127), dissolved in water to 10 mM stock, and used at 10 μM final concentration. Dopamine hydrochloride was purchased from Sigma Aldrich (Cat. #H8502), dissolved in 100 mM ascorbic acid to 10 mM stock, and used at 10 μM final concentration. DAMGO was purchased from Tocris (Cat. #1171), dissolved in DMSO to 10 mM stock, and used at 10 μM final concentration. 8-CPT-cAMP was purchased from Abcam (Cat. #ab120424), dissolved in water to 150 mM stock, and used at 150 μM final concentration for the RT-qPCR experiment. Forskolin was purchased from Sigma-Aldrich (Cat. #F6886), dissolved in DMSO to 10 mM stock, and used at 10 μM final concentration. Rolipram was purchased from Tocris (Cat. #0905), dissolved in ethanol to 10 mM stock, and used at 10 μM final concentration. Shield-1 ligand for stabilization of DD-tagged proteins was purchased from Aobious (Cat. #AOB1848), dissolved in ethanol to 1 mM stock, and added to the cell medium to 1 μM final concentration. D-luciferin sodium salt (Cat. #LUCNA) and coelenterazine (Cat. #CZ) were purchased from GoldBio and resuspended to 100 mM in 10 mM HEPES buffer, and 10 mM in ethanol, respectively, and stored protected from light. Dyngo-4a was purchased from Abcam (Cat. #ab120689), dissolved in DMSO to 30 mM stock, and added to cells grown in serum-free medium to a final concentration of 30 μM for 20 min prior to drug treatment.

### Constructs and siRNA

The previously described lentiCRISPRv2 vector for CRISPR KO gRNA expression was a gift from Feng Zhang (Addgene, Cat. #52961). gRNAs were cloned by annealing complementary oligonucleotides purchased from IDT (Table S1) and ligation into BsmBI-digested lentiCRISPRv2 as described previously.^61,62^ The parental vector for CRISPRi gRNA expression under a U6 promoter (pU6-gRNA-EF1alpha-puro-T2A-BFP) was a gift from Jonathan Weissman (Addgene, Cat. #60955).^63,64^ gRNAs were cloned by annealing complementary oligonucleotides purchased from IDT (Table S1) and ligation into BstXI/BlpI digested pU6-gRNA-EF1alpha-puro-T2A-BFP backbone as described previously.^8^

The plasmid encoding the cAMP luminescence biosensor with *Renilla* luciferase (pSF-CMV-GloSensor20F-IRES-Rluc, pGLO) and CMV promoter-driven FLAG-tagged-β2-AR were described previously.^8,13,17^ *DRD1*-Tango was a gift from Bryan Roth (University of North Carolina, NC) (Addgene, Cat. #66268). pcDNA 3.1-D2R, pcDNA 3.1-delta opioid receptor, and pcDNA 3.1-SSF-mu opioid receptor were a gift from Mark von Zastrow (University of California, San Francisco, CA) and transfected for 24h in HEK293 cells seeded on 6-well plates. Plasmids encoding TagBFP-PKAcat and ExRai-AKAR2 sensor were a gift from Jin Zhang (University of California, San Diego, CA) and transfected for 24h in HEK293 cells seeded on 6-well plates (TagBFP-PKAcat) or 35 mm imaging dishes (ExRai-AKAR2). The low expression β2-AR plasmid was generated by amplifying a genomic region that included the ~1 kb sequence upstream of the *ADRB2* ORF, which contains the annotated 5’ UTR and promoter, and the gene coding sequence. β2-AR and CMV-β2-AR plasmids were transfected for 48h in HEK293 cells seeded on 6-well plates.

To generate the EGFP-RBM12 plasmid, the wild-type human *RBM12* open reading frame sequence was PCR amplified from a plasmid encoding human *RBM12* in pDONR221 (DNASU, Cat. #HsCD00042134) and inserted into SacI-digested pEGFP-C1 backbone by In-Fusion cloning with an added stop codon. EGFP-c.2377G>T-RBM12 and EGFP-c.2532delT-RBM12 were generated using QuikChange site-directed mutagenesis of the wild-type EGFP-RBM12. To generate lentiviral plasmids encoding the wild-type and mutated EGFP-RBM12 constructs, the *RBM12* sequence was PCR amplified and inserted into XbaI-digested lentiviral FHUGW-EGFP (gift from Mark von Zastrow, University of California, San Francisco, CA) backbone by In-Fusion cloning.

A lentiviral plasmid encoding a transcriptional reporter for CREB activity (FHUGW-CRE-DD-zsGreen) was previously described.^8^ To generate a CREB activity reporter driving the production of the tdTomato fluorescent protein (FHUGW-CRE-DD-tdTomato), a sequence encoding the tdTomato fluorescent protein was amplified from a pBa-KIF5C 559-tdTomato-FKBP plasmid (Addgene, Cat. #64211) and inserted into the linearized FHUGW-CRE-DD-zsGreen plasmid by In-Fusion cloning.

Synthetic RNA duplexes (RBM12_8, Cat. #1027417; AllStar Negative Control, Cat. #1027281) were obtained from the validated HP GenomeWide siRNA collection (Qiagen) and transfected for 72 h using Lipofectamine-RNAiMax (Invitrogen, Cat. #13778150) according to the manufacturer’s instructions.

### Cell culture

HEK293 (from Mark von Zastrow, University of California, San Francisco, CA) and HEK293T (TakaraBio, Cat. #632180) cells were grown at 37°C/5% CO2 in Dulbecco’s Modified Eagle Medium (4.5 g/L glucose and L-glutamine, no sodium pyruvate) (Thermo Fisher Scientific, Cat. #11965118) supplemented with 10% fetal bovine serum (Sigma Aldrich, Cat. #F2442).

Human iPSCs engineered to express *NGN2* under a doxycycline-inducible system in the AAVS1 safe harbor locus were described previously.^32^ Human iPSCs stably-expressing FLAG-β2-AR were generated by lentiviral transduction, labeling with M1-Alexa-647 and fluorescent cell sorting. hiPSCs were cultured in Essential 8 Medium (Thermo Fisher Scientific, Cat. #A1517001) on plates coated with Growth Factor Reduced, Phenol Red-Free, LDEV-Free Matrigel Basement Membrane Matrix (Corning, Cat. #356231) diluted in Knockout DMEM to 0.1 mg/mL concentration (Thermo Fisher Scientific, Cat. #10829-018). Essential 8 Medium was replaced every 2 days. For lentiviral infection of hiPSCs, cells were infected for 3-4 days before neuronal differentiation.

### Lentivirus production

HEK293T cells were transfected with lentiviral constructs (wild-type or mutated EGFP-RBM12, CREB reporter, or CRISPRi gRNA) and standard packaging vectors (VSVG and psPAX2) using Lipofectamine-2000 (Invitrogen, Cat. #11668027) following recommended protocols. Supernatant was harvested 72 h after transfection and filtered through a 0.45 μm SFCA filter. The harvested virus was either used on the same day or concentrated and snap-frozen before use.

### HEK293 CRISPR KO and CRISPRi

RBM12 CRISPR knockout HEK293 clones were generated by transfecting wild-type cells with two independent gRNAs (Table S1) using Lipofectamine-2000 transfection reagent (Thermo Fisher, Cat. #11668027) following recommended protocols. Cells were selected with 1 μg/ml puromycin for 2 days, then recovered and plated as single colonies. Individual clones were expanded for at least 3 weeks and tested for successful editing using Sanger sequencing of genomic DNA. Parental wild-type cells were grown in parallel and serve as experimentally matched control. Sanger sequencing of the RBM12 KO cells shows the occurrence of insertion and deletion events that lead to a frameshift and a premature stop in both lines (Figure S2a). Using RT-qPCR analysis, we found that *RBM12* RNA levels were unaffected in KO 1 but reduced in KO 2 (Fig. S2b).

Wild-type and RBM12 KO HEK293 cells stably expressing FLAG-tagged β2-AR were generated by transfecting cells seeded on 6-well plate with the FLAG-β2-AR plasmid for 72 hours prior to selection of stable cells using 100 μg/mL G418 sulfate (Genesee Scientific, Cat. #25-538).

For CRISPRi-mediated knockdown, control gRNA (NTC) and *RBM12*-targeting gRNA (RBM12_783) were cloned into the parental vector for gRNA expression under a U6 promoter (pU6-gRNA-EF1alpha-puro-T2A-BFP) at the BlpI/BstXI sites. Previously described HEK293 cells stably expressing dCas9-BFP-KRAB^8^ were seeded on 6-well dishes at ~30% confluence and transduced with lentiviral supernatant containing the gRNAs of interest with or without the fluorescent CREB reporter. 48 h after transduction with the gRNAs, cells were selected with 1 μg/ml puromycin for 2 days, then recovered and expanded for 3 days in regular medium without antibiotic. For rescue experiments in HEK293, equal volumes of viral supernatant containing wild-type or mutated RBM12 lentiviral constructs were transduced together with the gRNAs on the first day. To generate stable CRISPRi-mediated *RBM12* knockdown hiPSCs, cells were transduced with lentiviral supernatant containing NTC or *RBM12*-targeting gRNA for 2 days followed by fluorescent cell sorting of gRNA-positive cells. For rescue experiments in neurons, NTC or *RBM12* gRNA-transduced hiPSCs were transduced with lentiviral supernatant containing EGFP vector, EGFP-WT, EGFP-c.2377G>T, or EGFP-c.2532delT RBM12 for 2 days and sorted based on EGFP expression prior to differentiation into neurons.

### iNeuron differentiation

iNeurons were generated by hiPSC differentiation for 14 days as described previously.^32^ Briefly, hiPSCs were lifted using Accutase and centrifuged. Pelleted cells were resuspended in N2 Pre-Differentiation Medium containing the following: Knockout DMEM/F12 (Thermo Fisher Scientific, Cat. #12660-012) as the base, 1X MEM Non-Essential Amino Acids (Thermo Fisher Scientific, Cat. #11140-050), 1X N2 Supplement (Thermo Fisher Scientific, Cat. #17502-048), 10ng/mL NT-3 (PeproTech, Cat. #450-03), 10ng/mL BDNF (PeproTech, Cat. #450-02), 1 μg/mL Mouse Laminin (Thermo Fisher Scientific, Cat. #23017-015), 10nM Y-27632 dihydrochloride ROCK inhibitor (Tocris, Cat. #125410), and 2 μg/mL doxycycline hydrochloride (Sigma-Aldrich, Cat. #D3447-500MG) to induce expression of *NGN2*. iPSCs were counted and plated at 5 × 10^5^ cells per Matrigel-coated well of a 6-well plate in N2 Pre-Differentiation Medium with 1 μg/mL doxycycline hydrochloride for 3 days. Afterwards, pre-differentiated cells were lifted and centrifuged as above, and the cell pellet was resuspended in Classic Neuronal Medium containing the following: half DMEM/F12 (Thermo Fisher Scientific, Cat. #11320-033) and half Neurobasal-A (Thermo Fisher Scientific, Cat. #10888-022) as the base, 1X MEM Non-Essential Amino Acids, 0.5X GlutaMAX Supplement (Thermo Fisher Scientific, Cat. #35050-061), 0.5X N2 Supplement (Thermo Fisher Scientific, Cat. #17502048), 0.5X B27 Supplement (Thermo Fisher Scientific, Cat. #17504-044), 10ng/mL NT-3 (PeproTech, Cat. #450-03), 10ng/mL BDNF (PeproTech, Cat. #450-02), 1 μg/mL Mouse Laminin (Thermo Fisher Scientific, Cat. #23017015), and 1 μg/mL doxycycline hydrochloride (Sigma-Aldrich, Cat. #D3447). Pre-differentiated cells were counted and plated at 3.5 × 10^5^ cells per well of a BioCoat Poly-D-Lysine 12-well plate (Corning, Cat. #356470) in Classic Neuronal Medium, or at 7 × 10^6^ cells per well of a BioCoat Poly-D-Lysine 6-well plate (Corning, Cat. #356469). After 7 days, half of the medium was removed and an equal volume of fresh Classic Neuronal Medium without doxycycline was added. After 14 days, half of the medium was removed and twice the volume of fresh Classic Neuronal Medium without doxycycline was added. For EGFP-RBM12 live-cell imaging experiments, iPSCs were counted and plated at 2 × 10^5^ cells per Matrigel-coated well of 35 mm imaging dish (Matsunami, Cat. #D1113OH) in N2 Pre-Differentiation Medium for 3 days, followed by media change using the Classic Neuronal Medium to induce differentiation.

### RT-qPCR analysis of target gene expression

For RT-qPCR experiments, cells were left untreated or treated with the indicated dose of drug for 1 hour in DMEM + 10% FBS prior to RNA extraction. For Dyngo-4a experiments, cells were pre-treated with DMSO or 30 μM Dyngo-4a in serum-free DMEM for 20 min prior to treatment with 1 μM Iso for 1 hour. Total RNA was extracted from the samples using the Zymo Quick-RNA Miniprep Kit (Genesee, Cat. #11–327) or Qiagen RNeasy Mini Kit (Qiagen, Cat. #74106). Reverse transcription was carried out using iScript RT supermix (Biorad, Cat. #1708841) or Superscript II Reverse Transcriptase (Thermo Fisher Scientific, Cat. #180644022) following recommended manufacturer protocols. The resulting cDNA was used as input for RT-qPCR using CFX-384 Touch Real-Time PCR System (Biorad), Power SYBR Green PCR MasterMix (Thermo Fisher Scientific, Cat. #4367659), and the appropriate primers (Table S2). All transcript levels were normalized to the levels of the housekeeping gene *GAPDH*.

### cAMP production

For pGLO sensor real-time measurement of cAMP production, cells seeded on 6-well plates were transfected with pGLO-20F/Rluc alone or with the indicated receptor plasmid (D1R, D2R, mu OR, delta OR) for 24 h using Lipofectamine 2000 transfection reagent following manufacturer’s protocols. On the day of the experiment, cells were replated onto a 96-well plate in medium supplemented with 160 μM D-luciferin and incubated for 40 min prior to conducting the assay. To measure cAMP production in response to Gαs-receptor activation, Hamamatsu FDSS/μCell with liquid handling was equilibrated at 37°C and used to dispense the drugs (100 nM isoproterenol for β2-AR, 10 μM NECA for A1/2R, 10 nM SKF-81297 for D1R) and simultaneously image cAMP-driven luciferase activity in real time. All experimental cAMP measurements (firefly luciferase time course data) were normalized to the *Renilla* luciferase signal (expression control), and the averaged normalized maximum values from the control samples for each tested batch was set to 100%, and all values are shown as % of this mean. D1R expression in transfected cells incubated with anti-HA-Alexa-488 (Thermo Fisher Scientific, Cat. #A-21287) for 1 hour on ice was measured by flow cytometry using BD FACS Canto2. To measure Gαi-receptor responses, cells were either treated with 1 μM forskolin + vehicle (DMSO), 1 μM forskolin + 10 μM DOPA + 10 μM ICI-118,551 (D2R), or 1 μM forskolin + 10 μM DAMGO (μOR and △OR). At the end of the time course, cells were lysed in stop buffer (5 mM HEPES, 2% glycerol, 1 mM EDTA, 400 uM DTT, 0.2% Triton) supplemented with 2 μM coelenterazine. All experimental cAMP measurements (firefly luciferase time course data) were normalized to the *Renilla* luciferase signal (expression control), and the inhibition of forskolin response (GPCR drug / forskolin) was calculated from each cell line. The averaged forskolin inhibition in the control samples for each tested batch was set to 100%, and all values are shown as % of this mean.

For the cADDis sensor, neurons were transduced with the BacMam sensor according to manufacturer’s instructions for 24 hours. On the day of the experiment, neurons were lifted with papain (Sigma-Aldrich, Cat. #P4762) and 100,000 cells were resuspended in 100 μL HBSS (Thermo Fisher Scientific, Cat. #14175-095) supplemented with 30 mM HEPES (Sigma Aldrich, Cat. #H0887) per well prior to drug addition and fluorescence reading using the TECAN Spark plate reader. For ELISA experiments, Cyclic AMP ELISA Kit (Cayman Chemical, Cat. #581001) was used according to manufacturers’ instructions and read using the TECAN Spark plate reader. All values were normalized to total protein amounts and shown as % of wild-type cells’ value.

### Receptor trafficking

Wild-type and RBM12 KO cells stably expressing FLAG-tagged β2-AR were seeded on 12-well plates. To induce β2-AR internalization, cells were treated with 1 μM isoproterenol for 20 min. Then, cells were put on ice to stop trafficking, lifted, and labeled with Alexa 647-conjugated M1 antibody. To induce β2-AR recycling, cells were first treated with 1 μM isoproterenol for 20 min and then 10 μM alprenolol for 40 m in. Untreated cells served as a proxy for total β2-AR cell surface expression. Flow cytometry analysis was carried out using a BD FACS Canto2 instrument, and Alexa-647 mean signal of the gated singlet population was used as a proxy for total number of surface receptors. Calculations were carried out for each cell line as follows: % internalized receptors = 100%-(total # surface receptors after 20 min isoproterenol)/(total # surface receptors pre-drug) × 100%; % recycled receptors = (total # surface receptors 40 min after alprenolol - total # surface receptors after 20 min iso) / (total initial # surface receptors - total # surface receptors after 20 min iso)] x 100%. The mean Alexa-647 wild-type values for the batch were set to 100%, and all values are shown as % of this mean.

### *Flow cytometry-based experiments with pCRE-DD-zsGreen1* and *pCRE-DD-tdTomato*

For fluorescent CREB reporter experiments, cells were seeded on 6-well plates and transduced with lentiviral supernatant containing the fluorescent CREB reporter.^8^ Cells were maintained for 5-7 days before experiments. On the experiment day, cells were treated for 4 hours with 1 μM Shield-1 ligand alone (basal) or with 1 μM Shield-1 ligand and one of the following: 1 μM isoproterenol, 10 μM epinephrine, 10μM norepinephrine, 50 nM salbuterol, 10 μM terbutaline, 10 μM formoterol prior to fluorescence reading using the BD FACS Canto2 flow cytometry instrument. For pCRE-DD-tdTomato experiments, cells were treated with 1 μM Shield-1 ligand alone (basal) or with 1 μM Shield-1 ligand and 1 μM isoproterenol for 6h. We used a BD LSRFortessa flow cytometry instrument and gated for gRNA-expressing (BFP+) and RBM12 construct-expressing (GFP+) singlets. From these measurements, the fold change (induced / basal) mean of the NTC gRNA + empty vector cells was averaged and set to 1, and the PE (tdTomato) mean for all other samples was normalized to this value (expressed as fold of mean NTC value).

### Microscopy

HEK293 cells were seeded on poly-L-lysine (Sigma, Cat. #P8920) coated coverslips in 12-well plates. iNeurons were plated on PDL-coated plates. Cells were fixed in 3.7% formaldehyde/PBS or 4% PFA/PBS for 20 min. Next, cells were stained using 1:100 – 1:500 anti-RBM12 primary antibody (Santa Cruz Biotechnology, Cat. #sc-514258) for 1-2 h and 1:1,000 Alexa fluor secondary antibodies for 30 minutes dissolved in 3% BSA/0.1% Triton-X/PBS blocking and permeabilizing solution. Lastly, cells were incubated with 1:5,000 DAPI (1 mg/ml stock) in PBS for 5 min.

For PKA activity measurements, HEK293 cells seeded on 35-mm imaging dishes (Matsunami, Cat. #D1113OH) were transfected with the ExRai-AKAR2 plasmid for 24 hours. On the day of the experiment, the medium was changed to Dulbecco’s Modified Eagle Medium (4.5 g/L glucose, no glutamine, no sodium pyruvate, no phenol red) (Thermo Fisher Scientific, Cat. #31053028) supplemented with 30 mM HEPES (Sigma Aldrich, Cat. #H0887), and cells were treated with the indicated drug concentration. Regions of interest (ROIs) were drawn around the cell to measure mean fluorescence values in FIJI. PKA activity was measured by calculating ΔF/F = (F_t_ - F_0_)/F_0_, with F_t_ representing the GFP fluorescence at a specific time point and F_0_ representing the initial GFP fluorescence.

Live and fixed cell imaging was performed on the Andor Dragonfly spinning disk microscope using the 40x/1.3 HC PL APO CS2 oil, WD: 0.24 mm objective lens (Leica, Cat. #11506358), 405-nm and 488-nm diode lasers (Andor), and 450/50 and 525/50 excitation/emission filters. The Andor iXon 888 Life EMCCD camera with 1024×1024 pixel was used with 200 EM gain. Images were collected with Andor Fusion Software. Live cell imaging was carried out in a 37°C chamber (Okolab).

### Protein extraction and Western blot

Cells were lysed with RIPA buffer (Sigma Aldrich, Cat. #R0278) containing protease inhibitors cocktail (Sigma-Aldrich, Cat. #P8340) and 0.1 μM PMSF (Sigma-Aldrich, Cat. #P7626). Lysates were then transferred to microcentrifuge tubes and spun for 5 min at 14,000 r.p.m. at 4 °C. The supernatant was used for western blot experiments. Protein concentration was determined by Pierce BSA Protein Assay Kit (Thermo Fisher Scientific, Cat. #23225). Protein samples were prepared for western blot analysis by adding 4x sample buffer (Bio-Rad, Cat. #1610747) and 1 mM dithiothreitol (Sigma, Cat. #D0632). Samples were loaded onto a 4-15% MINI-PROTEAN TGX Stain-Free gels (Biorad, Cat. #4568083) and transferred to nitrocellulose membrane for 1.5-3h at 100 V in 4 °C. Afterwards, membranes were blocked with 5% milk/TBST for 30 min and incubated with primary antibodies/TBST against proteins of interest: anti-RBM12 (Santa Cruz Biotechnology, Cat. #sc-514258), anti-PKA alpha polyclonal antibody (Thermo Fisher, Cat. #PA5-17626), and anti-beta-actin (Santa Cruz Biotechnology, Cat. #sc-69879). The following day, membranes were washed, incubated in the corresponding HRP-conjugated secondary antibodies/TBST and developed using Classico Western HRP substrate (Millipore Sigma, Cat. #WBLUC0100). Alternatively, membranes were washed, incubated with secondary antibody 1:10,000 diluted donkey anti-mouse-680 (LICOR Biosciences, Cat. #926-68072) and 1:10,000 diluted donkey anti-rabbit-800 (LICOR Biosciences, Cat. #926-32213) in LICOR blocking buffer (LICOR Biosciences, Cat. #927-40000), and visualized using the Odyssey imager system (LICOR). Bands were analyzed using densitometry analysis in ImageJ or the LICOR software. Scans of uncropped and unprocessed blots are provided.

### RNA-seq library preparation and analysis

RNA was extracted using the RNeasy Plus Mini Kit (Qiagen, Cat. #74106). 1 μg RNA and 500 ng RNA were used as input for sequencing libraries generation for HEK293 and neurons, respectively, and isolated using NEBNext Poly(A) mRNA magnetic isolation module (New England Biolabs, Cat. #E7490). HEK293 libraries were generated using NEBNext Ultra RNA Library Prep Kit for Illumina (New England Biolabs, Cat. #E7770) or NEBNext Ultra II Directional RNA Library Prep Kit for Illumina (#E7760), and NEBNext Multiplex Oligos for Illumina (New England Biolabs, Cat. #E7735S) according to the manufacturer’s instructions. Libraries were sequenced on NovaSeq 6000. Raw sequencing reads were analyzed using FastQC for quality control.^65^ Reads were aligned to the Homo sapiens GRCh38 assembly using STAR aligner.^66^ Raw read counts for each sample were obtained using featureCounts.^67^ Only genes with more than 5 counts in a minimum of 4 samples were included in the downstream analysis. Normalized counts and differential gene expression analysis was carried out in R using the DESeq2 pipeline.^68^

For the neuronal RNAseq analysis, we independently identified β2-AR target sets by differential expression analysis between untreated and isoproterenol-treated neurons in NTC and *RBM12* knockdown using DESeq2. Using a stringent statistical cut-off of gene p_adj_ < 5.0×10^-2^ (Wald test) and log_2_ fold change ≥ 0.5, we obtained 208 and 576 targets from NTC- and *RBM12* knockdown neurons, respectively. Of these, a total of 669 genes were unique and constituted the set of β2-AR-dependent transcriptional neuronal targets. For differential expression analysis between NTC and *RBM12* knockdown neurons, we applied a statistical cutoff of p_adj_ < 5.0×10^-2^ (Wald test) and log_2_ fold change ≥ 0.1 or ≤ 0.1. Gene Ontology analysis was performed using the two unranked list method in GOrilla^30^ using p_adj_ cutoff of < 5.0×10^-2^ (Fisher’s exact test).

Raw reads and normalized counts from the RNA-seq data generated in this study have been deposited on the Gene Expression Omnibus (GSE219195).

### Statistical analysis and reproducibility

All data are shown as mean ± SEM. Statistical analyses (included in source data files where applicable) to determine significance were performed using DESeq2 with Wald test for RNA-seq experiments and Prism v.8 (GraphPad) for unpaired t-test, one-, or two-way analysis of variance (ANOVA) (*α*, 0.05) for all other experiments. Asterisks are used to denote statistical significance (* = p ≤ 0.05, ** p ≤ 0.01, *** p ≤ 0.001, **** p ≤ 0.0001).

