## Supplementary Materials for "The psychosis risk factor *RBM12* encodes a novel repressor of GPCR/cAMP signal transduction"

#### Supplementary Figures

### Figure S1

**A**

| RBM12 KO #1 CRISPR gene editing | | | | | | | | | |
|---|---|---|---|---|---|---|---|---|---|
| WT | AGG | TCA | AGA | TCA | CCA | CAT | GAG | GCT | GGT |
| | $^{421}$R | S | R | S | P | H | E | A | G$^{429}$ |
| Edit 1 | AGG | TCA | AGA | TCA | CCA | CCA | TGA | GGC | TGG |
| | $^{421}$R | S | R | S | P | P$^{426}$ | STOP | | |
| Edit 2 | AGG | TCA | AGA | TGA | GGC | TGG | TTT | TTG | TGT |
| | $^{421}$R | S | R$^{423}$ | STOP | | | | | |

| RBM12 KO #2 CRISPR gene editing | | | | | | | | | | | | | | | | | |
|---|---|---|---|---|---|---|---|---|---|---|---|---|---|---|---|---|---|
| WT | AGG | GAA | ATG | ATA | CTA | AAT | CCA | GAG | GGG | GAT | GTC | AAC | TCT | GCC | AAA | GTC… | …ACA | AAG |
| | $^{529}$R | E | M | I | L | N | P | E | G | D | V | N | S | A | K | V | T | K$^{556}$ |
| Edit 1 | AGG | GAA | ATG | AGG | GGG | ATG | TCA | ACT | CTG | CCA | AAG | TCT | GTG | CCC | ACA | TAA | | |
| | $^{529}$R | E | M | R | G | M | S | T | L | P | K | S | V | P | T$^{543}$ | STOP | | |
| Edit 2 | AGG | GAA | ATG | TGC | CCA | CAT | AAC | AAA | TAT | TCC | ATT | CAG | CAT | TAC | AAA | GAT… | …GGA | TGA |
| | $^{529}$R | E | M | C | P | H | N | K | Y | S | I | Q | H | Y | K | D | G$^{556}$ | STOP |

**B**

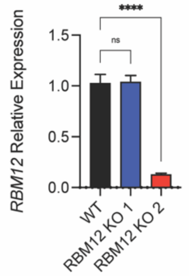


**C**

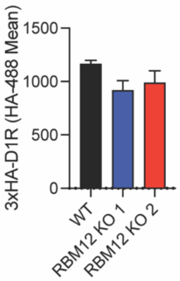


**D** **E** **F**

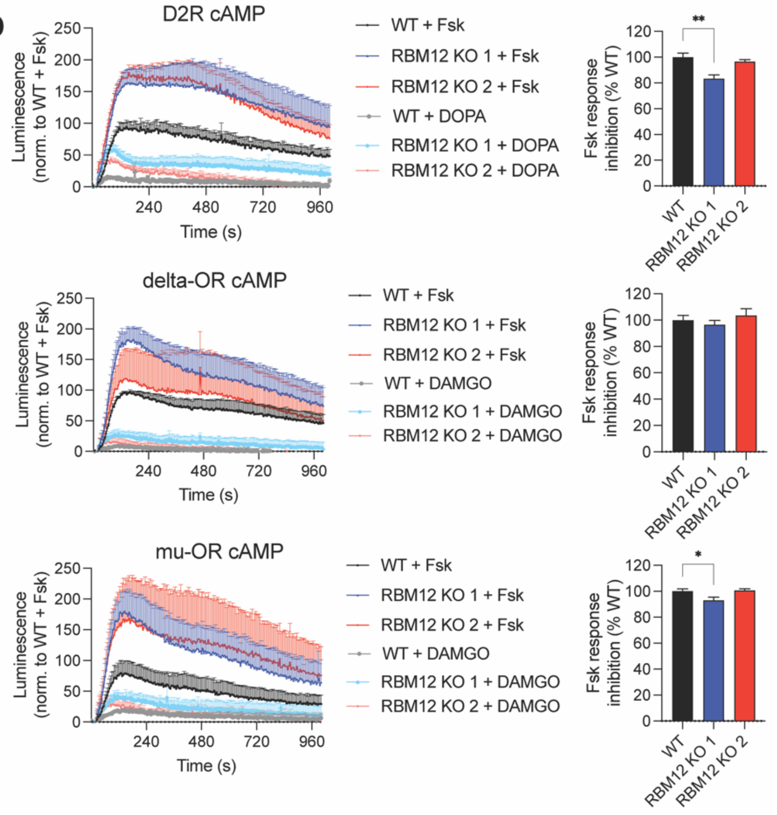

**Supplementary Figure 1. Characterization of RBM12 KO cell lines.** (A) Sanger sequencing of PCR-amplified genomic DNA from wild-type and RBM12 knockout clones showing positions of indels in each allele. (B) RT-qPCR of *RBM12* mRNA expression in the knockouts (n = 10). (C) Flow cytometry analysis showing comparable 3xHA-D1R expression in WT or RBM12 knockout cells transfected with plasmid encoding the receptor and surface-labeled with anti-HA-488 antibody (n = 4). (D-F) Luminescent GloSensor measurement of cAMP accumulation in cells overexpressing D2R (F, n = 4), ΔOR (G, n = 4), and μOR (H, n = 4) in response to either 10 μM forskolin and vehicle (DMSO) or 10 μM forskolin and 10 μM DOPA (D2R) in the presence of ICI-118,551 to isolate the D2R response (D), or 10 μM DAMGO (ΔOR and μOR) (E-F). All data are mean ± SEM. Statistical significance was determined using one-way ANOVA with Dunnett's correction (B, D-F).

**Figure S2**

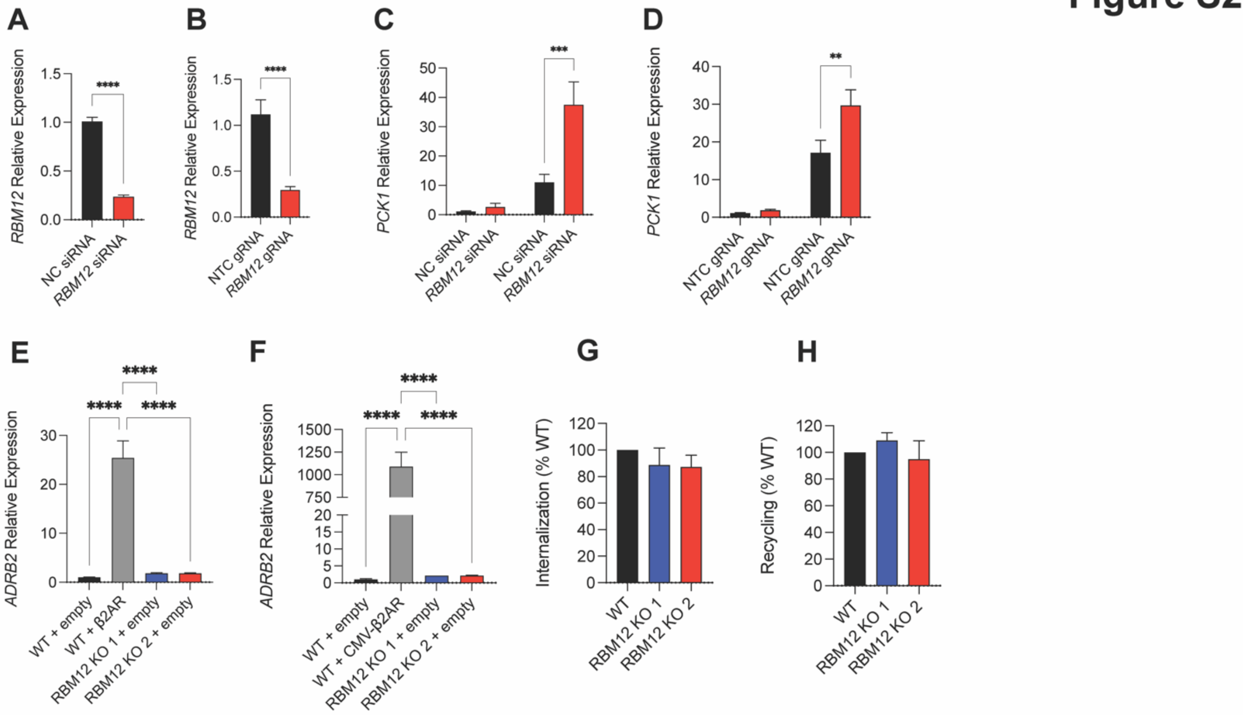


**Supplementary Figure 2. Knockdown efficiencies of different strategies to deplete RBM12 and characterization of β2AR overexpression.** (A-B) *RBM12* expression in cells transfected with non-targeting control (NC)- or *RBM12*-siRNA (A, n = 11), or NTC or *RBM12* CRISPRi gRNA (B, n = 10). (C) *PCK1* expression in cells transfected with non-targeting WT or *RBM12*-targeting siRNA, untreated or treated with 1 μM Iso for 1 hour (n = 13-14). (D) *PCK1* expression by RT-qPCR in cells expressing NTC or *RBM12*-targeting CRISPRi gRNA, untreated or treated with 1 μM Iso for 1 hour (n = 12). (E) *ADRB2* expression in cells transfected with empty plasmid or plasmid construct expressing β2AR from endogenous promoter (n = 3). (F) *ADRB2* expression in cells transfected with empty plasmid or plasmid construct expressing β2AR under a CMV promoter (n = 3). (G-H) Flow cytometry analysis of FLAG-β2AR internalization (G, n = 5) and recycling (H, n = 5). All data are mean ± SEM. Statistical significance was determined using unpaired t-test (A-B), two-way ANOVA with Tukey's correction (C-D), or one-way ANOVA with Tukey's correction.

### Figure S3

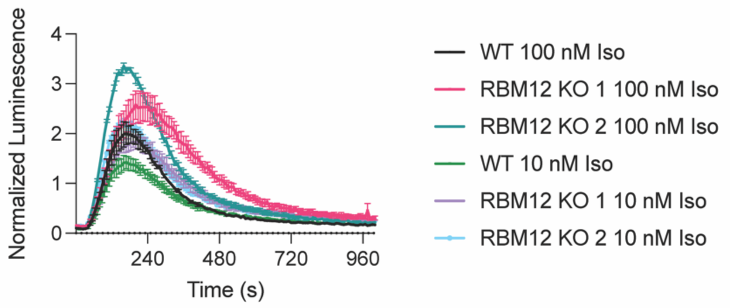


**Supplementary Figure 3. Isoproterenol dose-response measurement of cAMP production in wild-type and RBM12 KO cells.** Luminescent GloSensor measurement of cAMP accumulation following treatment with either 100 µM isoproterenol in wild-type cells or 10 nM isoproterenol in RBM12 knockout cells (n = 3). All data are mean ± SEM.

Figure S4

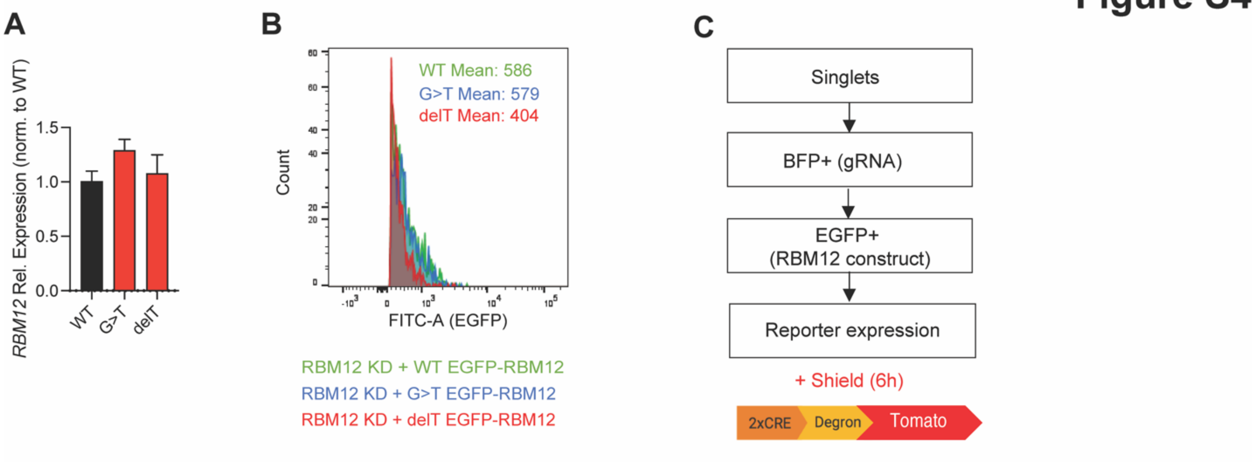


**Supplementary Figure 4. Wild-type and mutant RBM12 expression in HEK293 cells.** (A) Expression of *RBM12* mRNA by RT-qPCR in cells transfected with plasmid encoding EGFP-tagged WT, G>T, or delT RBM12 (n = 3). (B) Flow cytometry measurement of WT, G>T, or delT EGFP-RBM12 expression in the rescue assay. (C) Schematic of flow cytometry analysis strategy in the rescue assay.

Figure S5

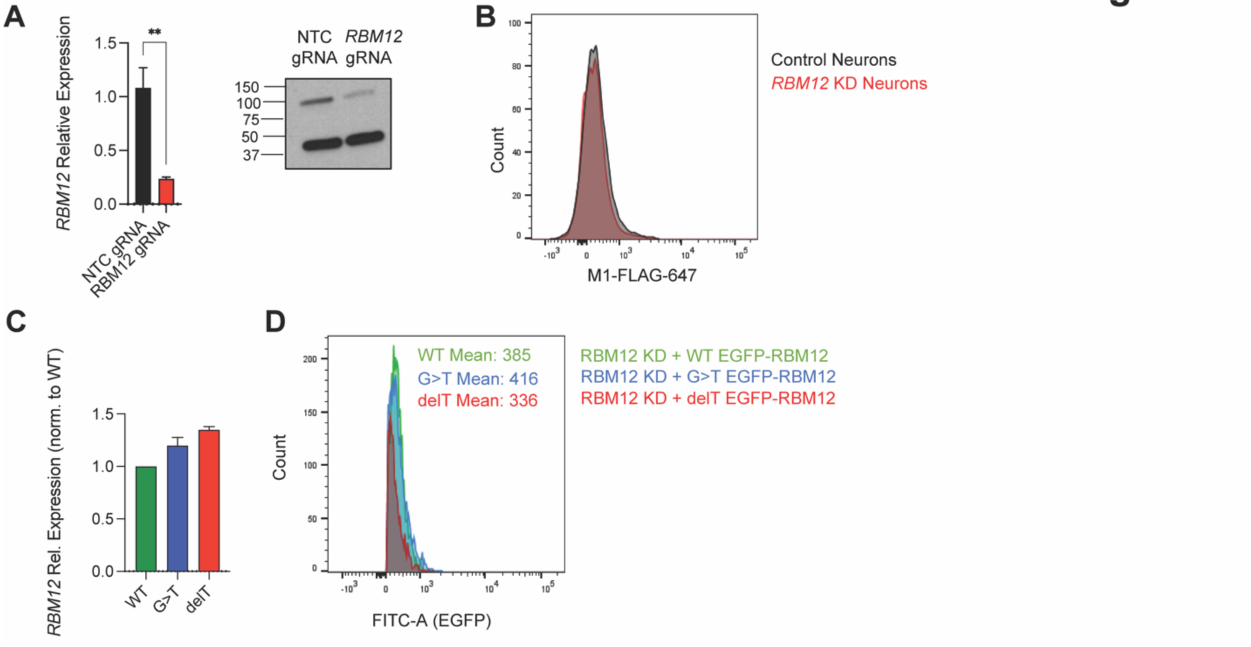


**Supplementary Figure 5. Generation and characterization of RBM12-depleted human neurons.** (A) *RBM12* expression (n = 6) and representative Western blot showing CRISPRi-dependent *RBM12* depletion in iNeurons. (B) Flow cytometry analysis of FLAG-tagged β2-AR expression in wild-type and *RBM12* KD neurons. (C) *RBM12* mRNA levels in neurons expressing WT, G>T, or delT EGFP-RBM12 in the rescue assay (n = 3). (D) Flow cytometry analysis of vector, WT, G>T, or delT EGFP-RBM12 expression in the neuron rescue assay. All data are mean ± SEM. Statistical significance was determined using unpaired t-test (A).

#### Figure S6

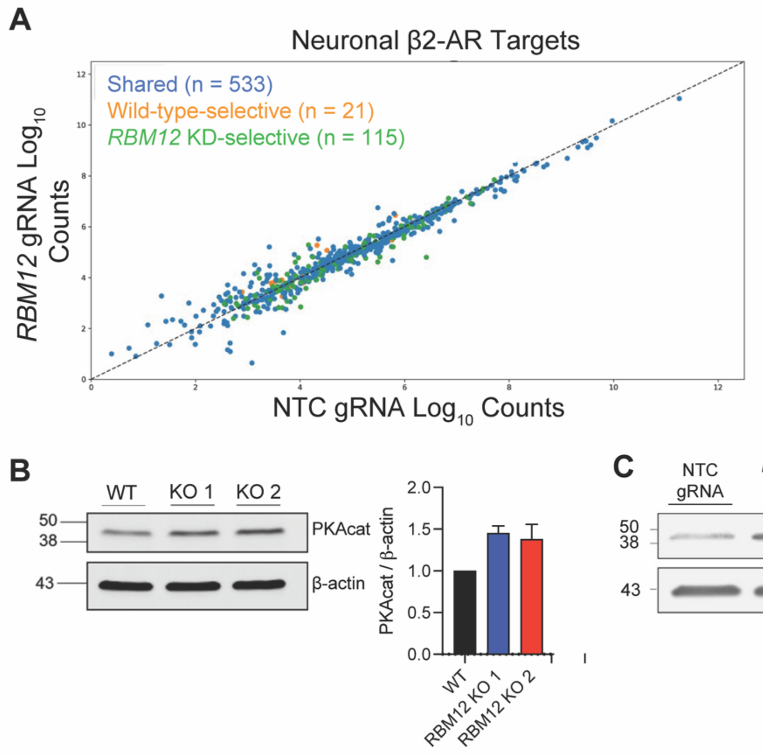


**Supplementary Figure 6. Basal expression of neuronal β2-AR target genes is unaffected by RBM12 depletion**. (A) Scatter plot of normalized RNAseq counts of neuronal β2-AR-dependent transcriptional targets in untreated cells (n = 669 genes). Blue dots represent genes that were induced by 1 hour 1 µM Iso treatment in both wild-type and *RBM12* KD neurons. Orange dots represent genes that were induced only in wild-type and unchanged or downregulated in *RBM12* KD neurons. Green dots represent genes that were induced only in *RBM1*2 KD neurons and unchanged or downregulated in wild-type. Indicated by arrows are a subset of genes with established roles in neuronal activity. The underlying information is summarized in Table 1. (B) Representative Western blot and quantification of the catalytic subunit of PKA (PKAcat) in wild-type and RBM12 knockout HEK293 cells, normalized to wild-type values per experiment (n = 2). (C) Representative Western blot and quantification of the catalytic subunit of PKA (PKAcat) in wild-type and *RBM12* knockdown neurons, normalized to wild-type values per experiment (n = 2).

#### Supplementary Tables

**Table S1. CRISPR KO and CRISPRi gRNA sequences used in this paper.**

| Name | Guide Sequence | Forward Primer (5' - 3') | Reverse Primer (5' - 3') |
|---|---|---|---|
| *RBM12* CRISPR KO gRNA #1 | AAGGTCAAGATCACCACATG | CACCGAAGGTCAAGATCACCACATG | AAACCATGTGGTGATCTTGACCTTC |
| *RBM12* CRISPR KO gRNA #2 | AAATGATACTAAATCCAGAG | CACCGAAATGATACTAAATCCAGAG | AAACCTCTGGATTTAGTATCATTTC |
| NTC CRISPRi gRNA | GGCAGGGCGTGGCGGGCGGTA | TTGGGCAGGGCGTGGCGGGCGGTAGTTTAAGAGC | TTAGCTCTTAAACTACCGCCCGCCACGCCCTGCCCAACAAG |
| *RBM12* CRISPRi gRNA | GAGGAGGTGGTGGCTGCGTT | TTGGAGGAGGTGGTGGCTGCGTTGTTTAAGAGC | TTAGCTCTTAAACAACGCAGCCACCACCTCCTCCAACAAG |

**Table S2. RT-qPCR primers used in this paper**

| Target Gene (*Homo sapiens*) | Forward Primer (5’-3’) | Reverse Primer (5’-3’) |
|---|---|---|
| *GAPDH* | CAATGACCCCTTCATTGACC | GACAAGCTTCCCGTTCTCAG |
| *PCK1* | CTGCCCAAGATCTTCCATGT | CAGCACCCTGGAGTTCTCTC |
| *NR4A1* | AGTGCAGAAAAACGCCAAGT | TTCGGACAACTTCCTTCACC |
| *FOS* | GCCTCTCTTACTACCACTCACC | AGATGGCAGTGACCGTGGGAAT |
| *RBM12* | GCCAAAGTCTGTGCCCACATAAC | GAACCAATGCCTGTCCTAGACC |
| *ADRB2* | GATTTCAGGATTGCCTTCCA | TATCCACTCTGCTCCCCTGT |
